# Targeting ZNRF3 and RNF43 to Restore Regeneration and Reverse Metabolic Dysfunction-Associated Steatotic Liver Disease

**DOI:** 10.1101/2025.06.19.660551

**Authors:** Federico Di Tullio, Sue Bin Yang, Lida Yang, Bruno Cogliati, Shamsa Roshan, Kaiyuan Guo, Abigail Glezer, Jonathan Conrad, Siddarth Vinod Kumar, Joana Almeida, Haoyuan Li, Shahina Saeed, Marina Barcena-varela, Emily Bramel, Anthony Lozano, Lianyong Su, Derrick Zhao, Huong Pham, Fanglin Ma, Amaia Lujambio, Sai Ma, Yizhou Dong, Huiping Zhou, Tianliang Sun

## Abstract

Liver regeneration and hepatocyte metabolic identity are disrupted in metabolic dysfunction-associated steatotic liver disease (MASLD) and its advanced form, metabolic dysfunction-associated steatohepatitis (MASH), yet the mechanisms of restore liver regeneration and reprogram metabolism for disease reversal remains poorly understood. Here, we show that β-Catenin activity progressively declines in hepatocytes during MASH in both humans and mice, coinciding with impaired regeneration and defective lipid clearance. Targeted deletion of the endogenous WNT pathway inhibitors ZNRF3 and RNF43 in hepatocytes after MASH onset reactivates β-Catenin signaling, leading to robust regression of steatosis, inflammation, and fibrosis, and restoring regenerative capacity across multiple fatty liver disease models. Mechanistically, this therapeutic effect is driven by β-Catenin–dependent induction of the alternative bile acid synthesis pathway without disrupting systemic lipid homeostasis. Importantly, both short-and long-term deletion of ZNRF3/RNF43 restores liver function without triggering tumorigenesis or hepatotoxicity, indicating a safe therapeutic window. These findings reveal that physiological activation of WNT/β-catenin signaling via ZNRF3 and RNF43 offers a viable regenerative and metabolic strategy for reversing fatty liver disease.

## Introduction

Fatty liver diseases including metabolic dysfunction-associated steatotic liver disease (MASLD) and its advanced version metabolic dysfunction-associated steatohepatitis (MASH) are major contributors to end-stage disease and hepatocellular carcinoma (HCC), affecting millions worldwide. Characterized by steatosis, inflammation, fibrosis, and impaired regeneration, MASH lacks broadly effective therapies due to its multifactorial signature. The inherited regenerative capacity allowing liver to deal with various injuries and precisely restore its mass and function without oncogenic risks. Thus, strategies by unlocking its regenerative power in a controlled way holds a promising potential for MASH therapy.

Elevated hepatic cholesterol serves as a key pathogenic driver, exacerbating hepatocyte injury and accelerates MASLD and MASH progression in both humans and animal models ^1–5^. Cholesterol clearance via bile acid synthesis— especially via the alternative pathway— represents a promising therapeutic approach^6–10^. Unlike the classical pathway, the alternative pathway generates chenodeoxycholic acid (CDCA)-enriched bile acids that are more efficient at cholesterol solubilization and hepatic lipid clearance. However, how to safely and effectively activate this pathway in fatty liver disease remains unresolved.

The WNT/β-Catenin pathway emerges as a unique therapeutic target that addresses both metabolic dysfunction and regenerative deficits in liver disease. This pathway plays a central role in regulating liver regeneration, metabolic homeostasis, and notably, bile acid synthesis—making it one of the few known signaling cascades capable of simultaneously enhancing cholesterol clearance and promoting hepatocyte proliferation. While β-catenin activation promotes hepatocyte proliferation in models of acute liver injury^11–13^, its role in chronic liver disease— especially in MASH—is more complex and remains incompletely understood. The pathway’s dual capacity to enhance regeneration and metabolic function positions it as an attractive target for MASH, yet concerns about potential risks such as tumorigenesis or cholestasis have limited therapeutic exploration.

The outcomes of WNT/β-catenin pathway activation are highly context-dependent, determined by three critical factors: (1) the method of pathway activation, (2) the underlying disease context, and (3) the duration and intensity of signaling. For instance, hepatocyte-specific deletion of Apc leads to strong β-catenin activation, promoting lipid clearance but causing bile acid accumulation and early lethality^14,15^. Conversely, gain-of-function mutations in β-Catenin result in hepatic lipid and bile acid accumulation with similarly reduced survival ^16,17^. These studies highlight the risks of excessive or dysregulated pathway activation. However, our previous work demonstrated that hepatocyte-specific deletion of ZNRF3 and RNF43—endogenous WNT inhibitors—enhances regeneration and reduces lipid accumulation under homeostasis conditions^18^. In contrast, other studies report that ZNRF3/RNF43 deletion under chronic injury can impair regeneration and promote lipid accumulation^19^, underscoring the importance of disease context. Crucially, while mutations in β-catenin pathway genes are common in HCC, β-catenin activation alone— particularly in the absence of other oncogenic stressors—is insufficient to initiate tumor formation. Different modes of pathway activation (e.g., ligand-dependent vs. mutation-driven) may result in distinct transcriptional outputs, with physiological activation promoting regenerative rather than tumorigenic programs^20^. Together, these findings underscore the importance of activation context and suggest that transient, physiological modulation of the WNT pathway—particularly through regulators such as ZNRF3/RNF43—may provide a therapeutic window to restore liver function and promote regeneration without triggering oncogenesis.

Here, we investigate the regenerative therapeutic potential of hepatocyte-specific deletion of ZNRF3 and RNF43 in two clinically relevant diet-induced MASH models that recapitulate the key features of human disease. We demonstrate that this intervention reverses fibrosis, steatosis, and inflammation while promoting robust hepatic regeneration without exacerbating liver injury or inducing HCC during the experimental period. Mechanistically, we show that β-catenin is required for MASH regression following ZNRF3/RNF43 deletion. This is mediated through selective activation of the alternative bile acid synthesis pathway, which promotes cholesterol conversion into CDCA-enriched bile acids and enhances both bile acid transport and lipid handling. Most importantly for translational applications, we demonstrate that transient ZNRF3/RNF43 knockdown using lipid nanoparticle (LNP)-delivered siRNAs —a platform already validated in FDA-approved therapeutics— induces MASH regression without tumor formation. These findings reveal a previously unrecognized regenerative mechanism driven by physiological WNT activation and establish ZNRF3/RNF43 as tractable therapeutic targets for reversing MASH through coordinated metabolic and regenerative reprogramming.

## Results

### Spatiotemporal dysregulation of canonical β-Catenin activity in hepatocytes during MASLD/MASH progression

WNT/β-Catenin signaling maintains liver metabolic zonation and regulates regeneration, yest its role in MASLD/MASH pathogenesis remains poorly understood^21^. To investigate cell type-specific β-Catenin activity across stages of MASLD, we analyzed a publicly available patient single-nucleus RNA sequencing (snRNA-seq) dataset ^22^. This dataset comprised liver biopsies from 47 patients across the MASH spectrum with detailed histopathological scoring. We focused on hepatocytes and assessed the relationship between β-Catenin target gene expression and disease severity. Unbiased clustering visualized by UMAP and scaled expression heatmaps of selected marker genes enabled identification of major liver cell types (**Suppl Figure 1A-B**). Among hepatocytes, expression of the canonical β-Catenin target gene *AXIN2* was elevated in early disease compared to healthy liver but progressively declined as disease advanced toward end-stage (**Figure 1A**). *AXIN2* expression also showed negative correlation with pathological features including steatosis, hepatocyte ballooning, liver inflammation and fibrosis (**Figure 1B**). Similar negative trends were observed for additional β-Catenin target genes in hepatocytes— including *CYP2E1, ZNRF3, and CYP27A1*—across multiple clinical features (**Suppl Figure 1C-E**).

**Figure 1.**
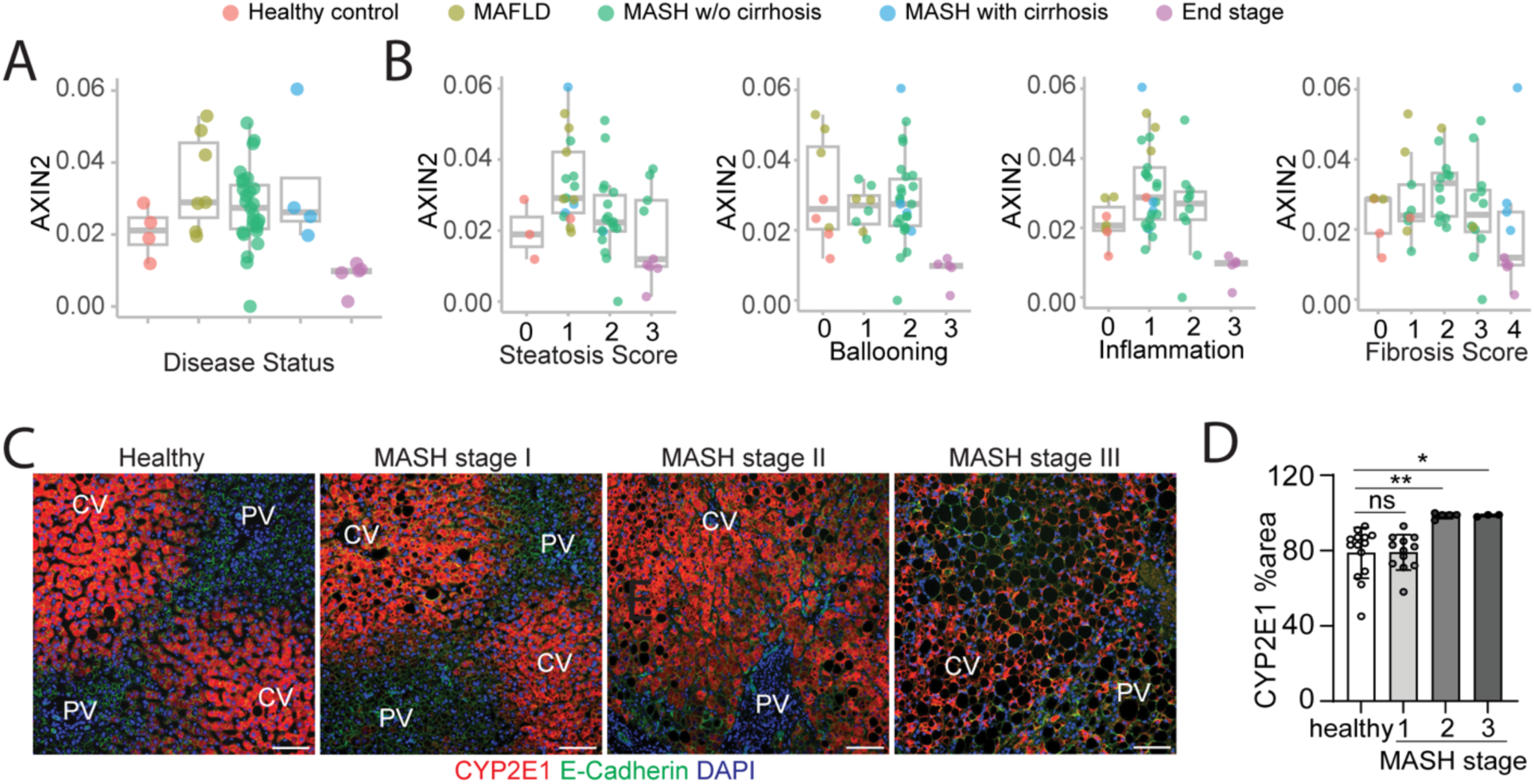
Impaired hepatocyte canonical β-Catenin activity and altered zonation across MASH progression. **(A)** AXIN2 expression in hepatocytes from human liver snRNA-seq data (GSE202379; n = 4 healthy, 7 MASLD, 27 MASH no cirrhosis, 4 MASH with cirrhosis, 5 end-stage). **(B)** AXIN2 expression stratified by histological features: steatosis, ballooning, inflammation, and fibrosis. **(C)** IF staining for CYP2E1 (red), E-cadherin (green), and DAPI (blue) in FFPE human liver sections (healthy, MASH stage I–III). **(D)** Quantification of CYP2E1-positive area (% area). CV, central vein; PV, portal vein. Data represent mean ± s.d.; *p < 0.05, **p < 0.01; ns, not significant. Two-tailed unpaired Student’s t-test. Scale bars: 100 µm (C).

Under healthy condition, β-catenin activity follows a pericentral-to-periportal gradient in hepatocytes that maintains metabolic zonation, with highest activity around central vein region.

Since previous studies reported breakdown of liver zonation during MASH progression^22^, we examined spatial WNT/β-Catenin activity in MASH livers across progression stages. Immunofluorescence staining of CYP2E1 in human MASH stage I livers showed a pericentral localization like healthy liver, but this signal progressively extended toward periportal hepatocytes in stages II and III, eventually involving nearly 100% of hepatocytes (**Figure 1C-D**).

Consistent with human data, CYP2E1 staining expanded from pericentral to periportal hepatocytes in the fibrosis and tumors (FAT)-MASH mouse model (Western diet combined with low-dose carbon tetrachloride)^5,23^ treated with 18 weeks compared to the chow diet controls (**Suppl Figure 1F-G**). Additionally, the reduction of β-Catenin target gene expression in hepatocytes was confirmed in the same MASH model by CYP1A2 western blot (**Suppl Figure 1H**).

These findings reveal that β-catenin activity becomes progressively suppressed in hepatocytes during MASLD/MASH progression, coinciding with metabolic dysfunction and impaired regeneration. Given that ZNRF3/RNF43 physiologically restrict β-catenin signaling through negative feedback, we hypothesized that their therapeutic deletion could restore β-catenin activity and reverse established MASLD/MASH pathology.

### ZNRF3/RNF43 deletion reverses established fatty liver diseases

To investigate the hepatocyte-specific role of ZNRF3/RNF43 in fatty liver diseases across distinct pathological contexts, we selected two complementary dietary models: the FAT-MASH model (Western diet + low-dose CCl_4_) for rapid fibrosis development^5,23^, and the Gubra Amylin (GAN) MASH diet model for closer recapitulation of human MASLD metabolic features^23^. We used adeno-associated virus type 8 (AAV8) vectors encoding Cre recombinase under the hepatocyte-specific TBG promoter (AAV8-TBG-Cre) to induce efficient hepatocyte-specific deletion of ZNRF3 and RNF43 in double-floxed mice (ZRNF3/RNF43^fl/fl^)^18^.

ZNRF3/RNF43^fl/fl^ mice were fed the FAT-MASH diet for 20 weeks to induce late-stage MASH characterized by inflammation, steatosis, and profound fibrosis, followed by intravenous injection of either AAV8-TBG-EGFP (control) or AAV8-TBG-Cre (ZNRF3/RNF43^ΔHep^). Mice remained on the Western diet for an additional 4 week before analysis (**Figure 2A, Suppl Figure 2A**). Compared to controls, ZNRF3/RNF43 deletion (ZRdKO) significantly reduced liver inflammation (H&E staining; **Figure 2B-C**), steatosis (Oil Red O staining; **Figure 2D-E**), and fibrosis (Sirius Red staining; **Figure 2F-G**). A single dose of AAV8-TBG-Cre efficiently induced knockout of ZNRF3 and RNF43 in hepatocytes, as evidenced by increased expression of the β-catenin target CYP2E1 throughout the liver (**Suppl Figure 2B-C**)^18,24^. Despite similar liver-to-body weight ratios (**Suppl Figure 2D**), ZNRF3/RNF43^ΔHep^ mice showed increased hepatocyte proliferation (Ki-67 staining; **Figure 2H-I**), indicating improved regenerative capacity, consistent with their previously reported role in response to acute injuries^18^. This unchanged liver-to-body weight ratio, despite increased proliferation, may be due to reduced hepatic lipid content offsetting the gain in cell mass.

**Figure 2.**
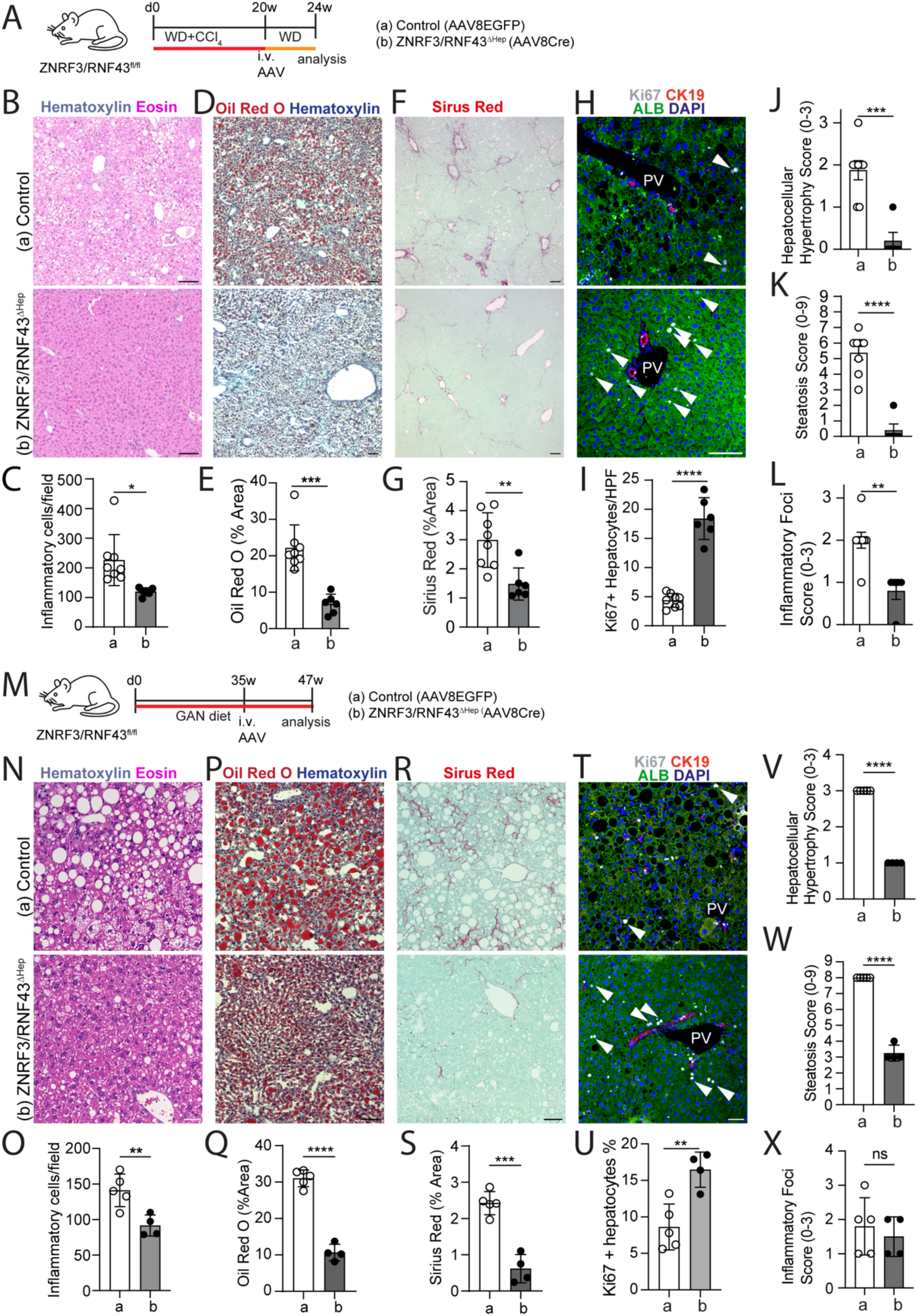
Hepatocyte-specific deletion of ZNRF3 and RNF43 promotes regeneration and reverses MASH pathology. Schematic of FAT-MASH model in ZNRF3/RNF43^fl/fl^ mice used for B-L. **(B–C)** H&E staining and quantification of inflammatory cell infiltration. **(D–E)** Oil Red O staining and quantification. **(F– G)** Sirius Red staining and quantification. **(H–I)** IF staining for Ki67 (gray), CK19 (red), Albumin (green), DAPI (blue) and quantification of Ki67⁺ hepatocytes. **(J–L)** Histopathological scoring of hypertrophy (**J)**, steatosis (**K)**, and inflammation (**L)**. **(M)** Schematic of GAN diet-induced model in ZNRF3/RNF43^fl/fl^ mice for N-X. **(N–O)** H&E staining and quantification of inflammatory cells. **(P–Q)** Oil Red O staining and quantification. **(R–S)** Sirius Red staining and quantification. **(T–U)** IF staining and Ki67⁺ hepatocyte quantification. **(V–X)** Histopathological scoring of hypertrophy (**V)**, steatosis (**W)**, and inflammation (**X)**. Data are presented as mean ± s.d., except for J-L and V-X, which are shown as mean ± s.e.m., with individual mouse values overlaid as dots. Two-tailed unpaired Student’s t-test. *p < 0.05; **p < 0.01; ***p < 0.001; ****p < 0.0001; ns, not significant. PV, portal vein. White arrowheads, Ki67⁺ hepatocytes. Scale bars: 100 µm (B, D, F, H, N, P, R, T).

Contrary to previous reports suggesting increased liver injury following ZNRF3/RNF43 deletion^19^, cleaved caspase-3–positive apoptotic hepatocytes (**Suppl Figure 2E-F**) and serum ALT levels (**Suppl Figure 2G**) were unchanged after 4 weeks of deletion. Histological improvements were confirmed by lower scores of hepatocellular hypertrophy (**Figure 2J**), steatosis with both macro-and microvesicular steatosis (**Figure 2K, Suppl Figure 2H-I**), and inflammation (**Figure 2L**) using a rodent MASH scoring system^25^.

To validate these findings and evaluate long-term regression, we employed the GAN diet MASH model. This model has recently been ranked as the most reflective of human MASLD features among commonly used murine dietary models^23^. ZNRF3/RNF43^fl/fl^ mice fed GAN diet for 35 weeks were injected with AAV8-TBG-EGFP or AAV8-TBG-Cre, followed by 12 weeks of continued GAN diet (**Figure 2M, Suppl Figure 2J**). Consistent with the FAT-MASH model, ZNRF3/RNF43 deletion reversed the GAN diet-induced MASH features, with significant reductions in immune cell infiltration (**Figure 2N-O**), steatosis (**Figure 2P-Q**), and fibrosis (**Figure 2R-S**). The efficient ZNRF3 and RNF43 knockout in hepatocyte was verified by the upregulation of CYP2E1 across the liver (**Suppl Figure 2K-L**). Liver-to-body weight ratios remained unchanged upon 12 weeks of ZNRF3/RNF43 knockout (**Suppl Figure 2M**), while increased hepatocyte proliferation was observed (**Figure 2T-U**). Notably, this was accompanied by a marked restore the loss of both the number and stacked morphology of endoplasmic reticulum (ER) structures in ZNRF3/RNF43-deficient hepatocytes just two weeks after gene deletion in mice previously maintained on the GAN diet for 28 weeks, suggesting early activation of a regenerative and metabolic remodeling program **(Suppl Figure 2S-T).**

While serum ALT levels remained unchanged after 4 weeks of ZRdKO in the FAT-MASH model (**Suppl Figure 2G**), serum ALT was reduced after 12 weeks of knockout in the GAN diet model (**Suppl Figure 2P**), indicating improved regeneration with decreased liver injury upon the longer treatment. Overall low cell apoptosis was detected by the cleaved caspase-3 staining in both groups, further supported no increased liver injury upon ZRdKO (Suppl Figure 2N-O). Rodent MASH score confirmed the improvement of histological features of hepatocytes by reduced hepatocellular hypertrophy (**Figure 2V**) and steatosis (**Figure 2W, Suppl Figure 2Q-R**). Of note, although inflammatory foci scores remained unchanged (**Figure 2X**), we observed a significantly reduced number of large immune cell clusters around residual steatotic hepatocytes in the ZRdKO group, resulting in an overall reduction in immune cell infiltration. Comprehensive histopathological examination by a blinded veterinary pathologist revealed no tumors, necrosis, or dysplastic nodules in any treated animals across both models and timepoints.

Collectively, hepatocyte-specific ZNRF3/RNF43 deletion consistently reversed key MASLD/MASH features across two complementary disease models, achieving therapeutic regression without hepatotoxicity or tumor formation. The robust nature of this therapeutic effect prompted investigation of whether β-catenin signaling—the canonical target of ZNRF3/RNF43— is required for MASH regression.

### β-Catenin dependency of reversing fatty liver disease induced by ZNRF3/RNF43 deletion

ZNRF3/RNF43 are E3 ubiquitin ligases that restrict WNT/β-Catenin activity by degrading Fizzled receptors^26,27^. To test whether β-Catenin is required for MASLD/MASH regression induced by ZNRF3/RNF43 deletion, mice were fed GAN diet for 16 weeks, followed by AAV8-mediated deletion of ZNRF3/RNR43 alone or together with a β-Catenin (CTNNB1), and continued GAN diet for four more weeks. To rigorously control for Cre recombinase and AAV8 infection effects, we included two additional control groups: (a) Rosa26-LSL-EGFP mice injected with AAV8-TBG-Cre (Cre-only control), and (b) ZNRF3/RNF43^fl/fl^ mice injected with AAV8-TBG-EGFP (virus-only control). Experimental groups were designed to assess the requirement for β-catenin in ZNRF3/RNF43-mediated MASH regression, including ZNRF3/RNF43 deletion alone (group c: ZNRF3/RNF43^τιHEP^ + AAV8Cre) and triple deletion of ZNRF3/RNF43/CTNNB1 in hepatocytes (group d: ZNRF3/RNF43/CTNNB1^τιHEP^ + AAV8Cre) (**Figure 3A, Suppl Figure 3A**).

**Figure 3.**
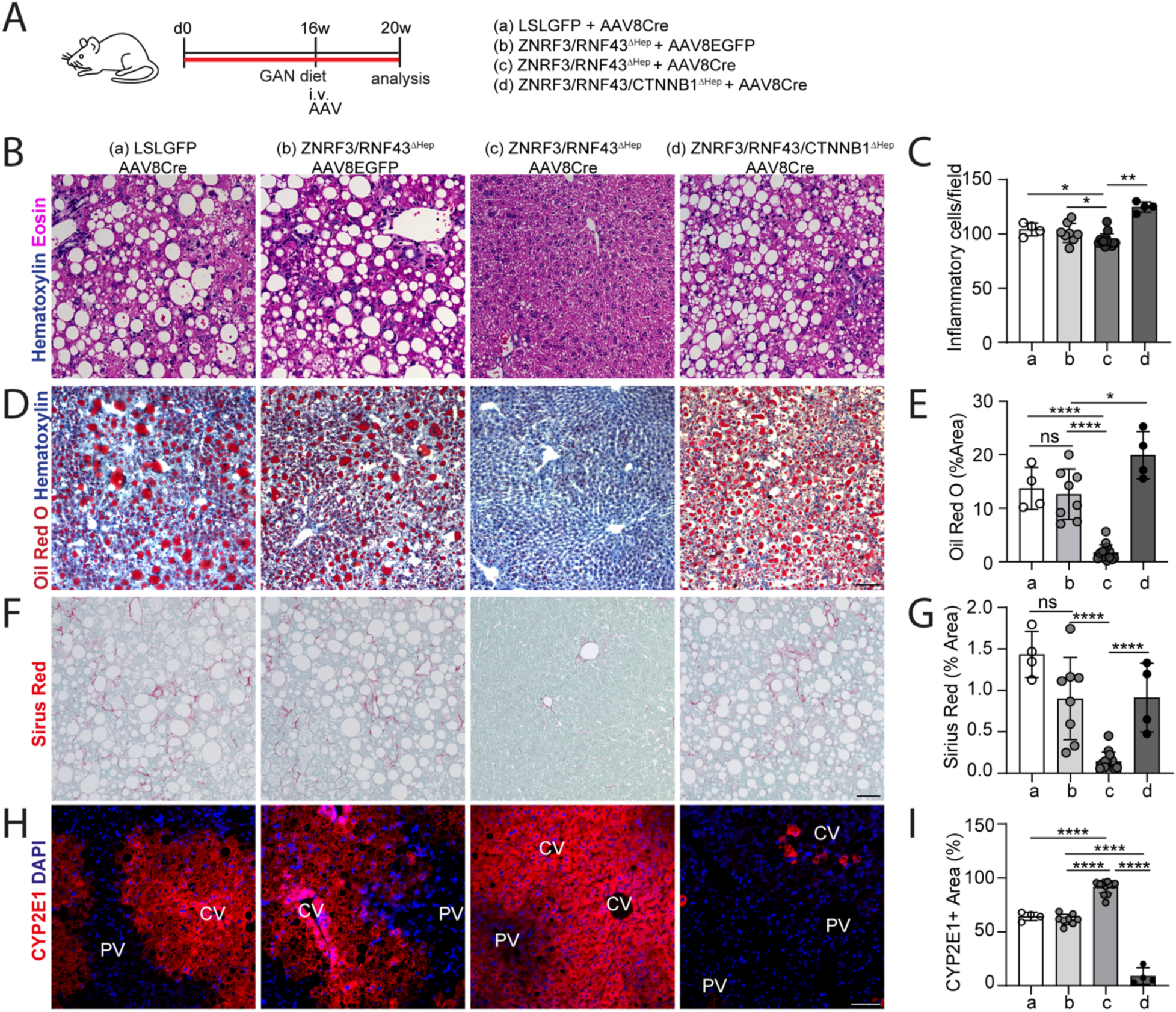
β-Catenin is required for mash regression following ZNRF3/RNF43 deletion. Schematic of the GAN diet-induced MASH model in four groups of mice for B-I. **(B–C)** H&E staining and inflammatory cell quantification. **(D–E)** Oil Red O staining and quantification. **(F–G)** Sirius Red staining and quantification. **(H–I)** IF for CYP2E1 (red) and DAPI (blue) and quantification of CYP2E1⁺ area. Data are presented as mean ± s.d., with individual mouse values overlaid as dots. Two-tailed unpaired Student’s t-test. *p < 0.05; **p < 0.01; ****p < 0.0001; ns, not significant. CV, central vein; PV, portal vein. Scale bars: 100 µm (B, D, F, H).

The two control groups (a and b) displayed comparable levels of inflammation (**Figure 3B-C, groups a and b**), steatosis (**Figure 3D-E, groups a and b**) and fibrosis (**Figure 3F-G, groups a and b**) while ZNRF3/RNF43 deletion alone significantly reduced these parameters as previously showed in Figure2 (**Figure 3B-G, group c**). Strikingly, simultaneous deletion of β-Catenin fully abolished the MASH regression induced by ZNRF3/RNF43 loss (**Figure 3B-G, group d**), indicating β-catenin is essential for the observed therapeutic effect. Efficient gene deletion was confirmed by immunostaining of the β-catenin target CYP2E1: ZNRF3/RNF43 deletion expanded CYP2E1 expression throughout the liver, while triple knockout nearly abolished its expression (**Figure 3H-I**). Consistent with 4 weeks regression in FAT-MASH model, liver to body ratio and serum ALT levels are not changed (**Suppl Figure 3B-C**). ZNRF3/RNF43 deletion did not alter systemic energy balance, as food intake and body weight gain remained unchanged (**Suppl Figure 3D-E**). In contrast, triple knockout mice exhibited reduced body weight gain, suggesting a hepatocyte-intrinsic role for β-catenin in maintaining energy homeostasis (**Suppl Figure 3D**). Altogether, these findings demonstrate that the therapeutic effects of ZNRF3/RNF43 deletion on MASLD/MASH regression—including reductions in steatosis, inflammation, and fibrosis—require β-catenin activity in hepatocytes.

### ZNRF3/RNF43 deletion activates cholesterol to bile acid conversion by selectively targeting the alternative bile acid synthesis pathway

Having established that β-catenin is essential for ZNRF3/RNF43-mediated MASLD/MASH regression, we next investigated the molecular mechanisms underlying this therapeutic effect, focusing on metabolic reprogramming pathways that could explain the coordinated improvements in steatosis, inflammation, and fibrosis. We first tested whether hepatic lipids are redistributed systemically, we monitored serum cholesterol and triglycerides levels following ZNRF3/RNF43 deletion in both GAN and FAT-MASH models. These levels remained unchanged (**Suppl Figure 3F-J**), ruling out systemic redistribution to circulation. This was confirmed by biweekly monitoring of serum cholesterol and triglycerides during 12 weeks of regression in GAN-fed mice (**Suppl Figure 3K-M**). Despite stable serum lipids, fecal cholesterol instead of triglycerides was significantly reduced after 12 weeks of ZRdKO (**Suppl Figure 3N-O**), suggesting the potential of hepatic depletion of cholesterol. The absence of ectopic lipid deposition in peripheral tissues including heart, kidney, and intestine supported a liver-intrinsic lipid disposal mechanism (**Suppl Figure 3P-U**). We next assessed hepatic lipid content. Triglyceride, total cholesterol, and free cholesterol levels were all significantly decreased in livers after 12 weeks of ZNRF3/RNF43 deletion (**Suppl Figure 4A-D**), confirming active lipid clearance from the liver.

To understand the underlying programs, we performed bulk RNA-seq on livers from FAT-MASH mice (20 weeks on diet) after 4 weeks of ZNRF3/RNF43 deletion (**Figure 4A, Suppl Figure 4E**). Gene set enrichment analysis (GSEA) revealed activation of metabolic pathways related to xenobiotic detoxification, fatty acid and cholesterol metabolism, bile acid synthesis, and oxidative metabolism (**Figure 4B, 4D**), alongside suppression of inflammatory, fibrotic, and apoptotic gene signatures (**Figure 4C, 4D**). These changes were corroborated by qPCR in a separate GAN model (**Suppl Figure 4G-I**).

**Figure 4.**
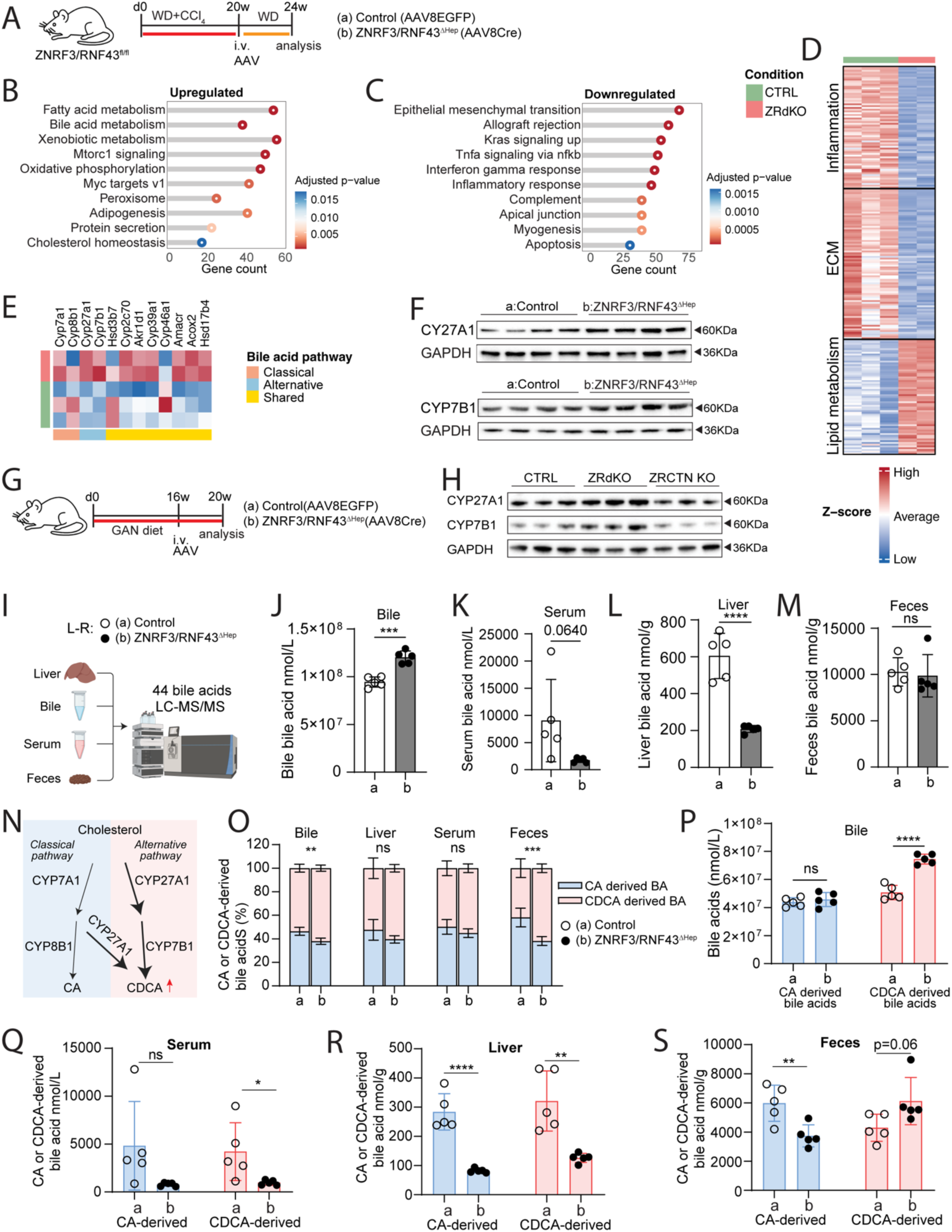
hepatocyte-specific ZNRF3/RNF43 deletion activates the alternative bile acid synthesis pathway. **(A)** Schematic of FAT-MASH model in ZNRF3/RNF43^fl/fl^ mice used for B-F. **(B–C)** GSEA of upregulated (B) and downregulated (C) pathways. **(D)** Heatmap of inflammatory, ECM, and lipid metabolism gene expression (z-score). **(E)** Heatmap of bile acid pathway genes categorized into classical (orange), alternative (turquoise), or shared (yellow) pathway. **(F)** Western blot of CYP27A1 and CYP7B1. **(G)** Schematic of GAN diet-induced model in ZNRF3/RNF43^fl/fl^ mice used for H-S. **(H)** Western blot of CYP27A1 and CYP7B1 from control, ZRdKO, and ZRCTN triple KO livers. **(I)** Schematic of LC-MS/MS bile acid profiling in liver, bile, serum, and feces. **(J–M)** Total bile acid concentrations in bile (J), serum (K), liver (L), and feces (M). **(N)** Schematic of bile acid biosynthesis pathways. **(O)** Stacked bar plots showing proportions of CA-derived and CDCA-derived bile acids. **(P–S)** Total concentrations of CA-derived and CDCA-derived bile acids in bile (P), serum (Q), liver (R), and feces (S). Data are presented as mean ± s.d., with individual mouse values overlaid as dots. Two-tailed unpaired Student’s t-test. *p < 0.05; **p < 0.01; ***p < 0.001; ****p < 0.0001; ns, not significant.

Interestingly, while lipid metabolic genes were broadly induced (**Figure 4D**), genes regulating lipogenesis (e.g., *Srebp2, Pparg, Cd36, Scd1*) and lipolysis (e.g., *Hsl, Cpt1a, Atgl*) were unchanged from FAT-MASH RNA-seq and qPCR in GAN diet MASH model (**Suppl Figure 4E-F, 4G, 4J-K**), indicating non-canonical lipid catabolism. RNA-seq also showed elevated expression of cholesterol biosynthetic and bile acid metabolic genes (**Suppl Figure 4E-F**), despite reduced hepatic and fecal cholesterol, prompting investigation into cholesterol disposal routes.

Given the central role of bile acid synthesis in cholesterol clearance, we examined pathway-specific gene expression. Remarkably, ZNRF3/RNF43 deletion selectively upregulated key genes in the alternative pathway (*Cyp27a1, Cyp7b1*) and bile acid transporters (*Abcg5, Abcg8, Abcb11, Slc10a1*), while leaving the classical pathway genes (*Cyp7a1, Cyp8b1*) unchanged (**Figure 4E**). These transcriptomic shifts were confirmed by western blot, which showed increased hepatic CYP27A1 and CYP7B1 protein levels in ZRdKO livers (**Figure 4F**).

To validate these findings in an independent model, we examined 16-week GAN-fed mice treated with ZNRF3/RNF43 deletion (**Figure 4G**). Both mRNA and protein levels of *Cyp27a1* and *Cyp7b1* from the alternative pathway were elevated, while classical pathway genes like *Cyp7a1* and *Cyp8b1* remained unchanged (**Figure 4H, Suppl Figure 4L-M**). Triple knockout of ZNRF3/RNF43/CTNNB1 abrogated this induction, confirming β-Catenin dependency (**Figure 4H**). RNA-seq analysis revealed selective upregulation of alternative bile acid synthesis genes, suggesting a novel mechanism for cholesterol clearance. To functionally assess bile acid synthesis and distribution, we profiled 44 major bile acids across liver, bile, serum, and feces in ZNRF3/RNF43-deleted and control mice via liquid chromatography tandem mass spectrometry (LC-MS/MS) (**Figure 4G, 4I**). Total bile acid levels were increased in bile but decreased in liver and serum, while fecal levels remained unchanged (**Figure 4J-M**), indicating efficient hepatic bile acid export without accumulation.

The classical bile acid synthesis pathway produces both cholic acid (CA) and chenodeoxycholic acid (CDCA), while the alternative pathway generates only CDCA. Therefore, we inferred the relative activity of each pathway by grouping detected bile acids into CA-derived (CA-BAs) and CDCA-derived (CDCA-BAs) species (**Figure 4N**). We then examined their ratio across tissues to evaluate the relative contribution of each synthesis pathway. The ratio of CDCA-to CA-derived bile acids was significantly increased in bile and feces, but unchanged in liver and serum (**Figure 4O**), indicating a shift toward CDCA-derived alternative bile acid synthesis. Absolute quantification confirmed a selective increase in CDCA-derived bile acids in bile, while CA-derived species remained stable, confirming the selective elevating of bile acid synthesis via the alternative pathway (**Figure 4P**). This compositional shift was echoed in feces, with elevated CDCA-derived and reduced CA-derived bile acids (**Figure 4S**). In contrast, both classes declined in liver and serum (**Figure 4Q-R**), suggesting altered enterohepatic cycling and enhanced biliary excretion. Importantly, this protective bile acid handling pattern—favoring CDCA derivatives without systemic overload—distinguishes ZNRF3/RNF43 deletion from other β-Catenin activation models (e.g., APC deletion and β-Catenin gain-of-function mutations), which often lead to bile acid accumulation and cholestatic liver diseases^15,16^.

This selectivity is therapeutically significant because the alternative pathway generates CDCA-enriched bile acids that are more efficient at cholesterol solubilization than classical pathway and less likely to cause cholestasis. Prior studies support the therapeutic benefits of boosting the alternative pathway to improve lipid metabolism and reduce inflammation and fibrosis ^6–10^.

### ZNRF3/RNF43 deletion reprograms hepatic bile acid composition toward a detoxified profile during MASLD/MASH regression

To determine whether the observed shift toward alternative bile acid synthesis translates into a therapeutically beneficial bile acid profile, we comprehensively analyzed bile acid composition across bile, serum, liver, and feces. Secondary bile acids —known to engage metabolic and immune signaling pathways via receptors such as FXR and TGR5—increased in bile (from 7% to 14%) and serum (from 11% to 30%), while remaining stable in liver and feces (**Figure 5A**). This bile shift was primarily driven by a fold increase in secondary species, with primary bile acids still predominant (**Figure 5B**). In contrast, shift in serum and liver bile acid levels were due to reduced primary species (**Figure 5C–D**), while fecal bile acids remained unchanged (**Figure 5E**).

**Figure 5.**
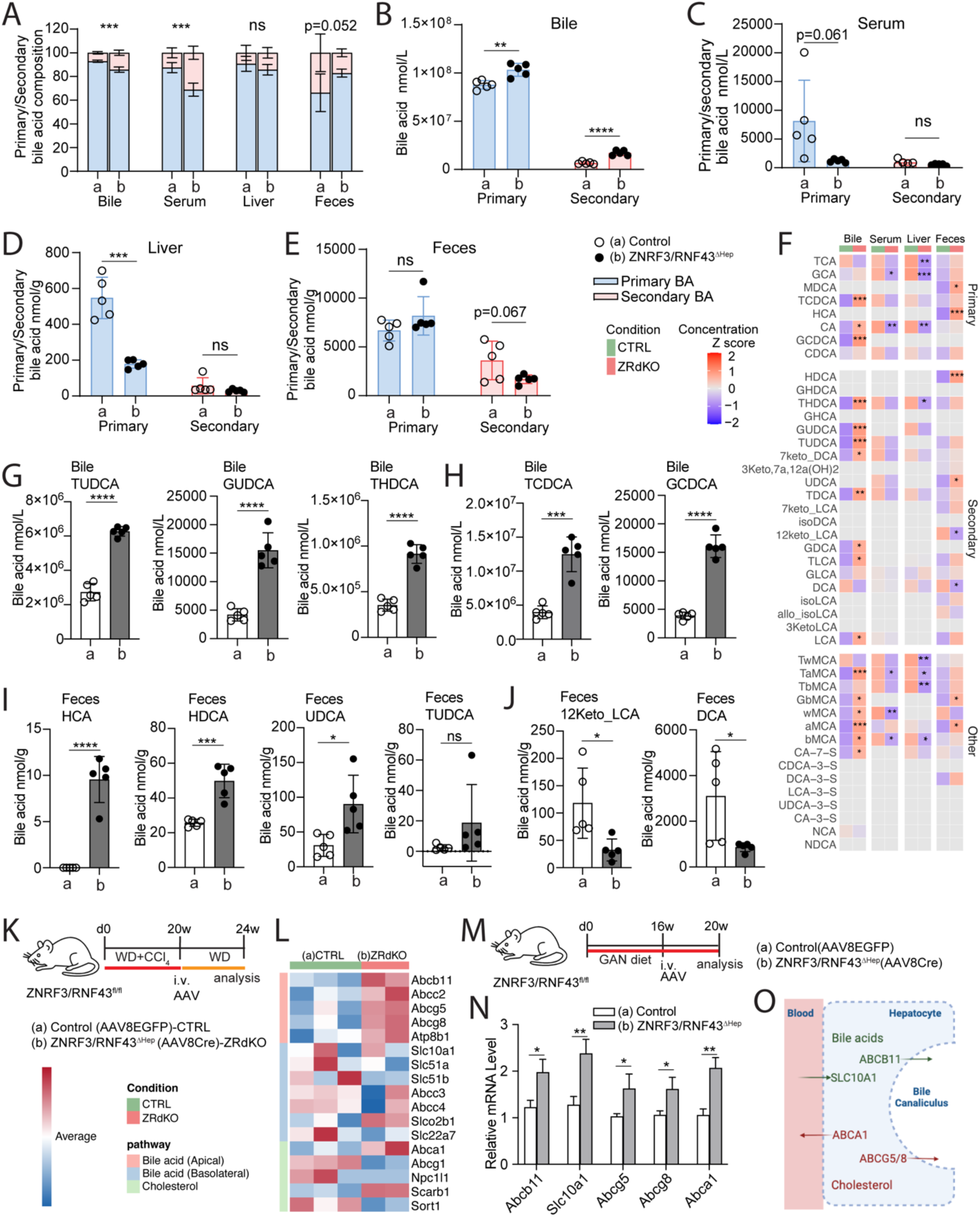
ZNRF3/RNF43 deletion reshapes bile acid composition and enhances lipid handling. Experimental design as described in Figure 4G for A-I. **(A)** Stacked bar plots showing primary (light blue) vs secondary (light pink) bile acids in bile, liver, serum, and feces. Groups: (a) Control, (b) ZNRF3/RNF43^Δhep^. **(B–E)** Total primary vs secondary bile acid concentrations in bile (B), serum (C), liver (D), and feces (E). **(F)** Heatmap of 44 bile acid species across tissues; rows grouped by bile acid class, columns by tissue/condition. **(G–H)** Concentrations of TUDCA, GUDCA, THDCA (G), and TCDCA, GCDCA (H) in bile. **(I–J)** Concentrations of HCA, HDCA, UDCA, TUDCA **(I)**, and 12Keto-LCA, DCA **(J)** in feces. **(K)** Schematic of FAT-MASH model in ZNRF3/RNF43^fl/fl^ mice used for L. **(L)** Heatmap of bile acid and cholesterol transporter expression; color-coded by transporter class. **(M)** Schematic of GAN diet-induced model in ZNRF3/RNF43^fl/fl^ mice used for N. **(N)** qPCR analysis of *Abcb11* (BSEP), *Slc10a1* (NTCP), *Abcg5*, *Abcg8*, and *Abca1*. **(O)** Schematic illustration of hepatocyte transporter localization from N. Data are presented as mean ± s.d., except for N, which are shown as mean ± s.e.m., with individual mouse values overlaid as dots. Two-tailed unpaired Student’s t-test. *p < 0.05; **p < 0.01; ***p < 0.001; ****p < 0.0001; ns, not significant.

Bile acid conjugation patterns were largely preserved. In liver and feces, conjugation ratios were stable (**Suppl. Figure 5A**). In bile and serum, the proportion of unconjugated bile acids increased slightly, but conjugated forms still dominated (**Suppl. Figure 5A**). ZRdKO induced increase of both conjugated and unconjugated bile acids in bile, while no change was observed in feces (**Suppl Figure 5B, 5E**). Interesting, both conjugated or unconjugated bile acids are reduced in serum and liver with conjugated bile acid dominated the change, suggesting the selective clearance of conjugated bile acids in these tissues (**Suppl Figure 5C-D**). Despite these changes, key conjugation genes (*Slc27a5, Baat*) were unaltered (**Suppl. Figure 5F**), suggesting preserved detoxification machinery.

Compared to the broader bile acid compositional data **(Figure 4J–M)**, the heatmap (**Figure 5F**) reveals an integrated view of individual bile acid shifts across tissues, highlights a coordinated enrichment of cytoprotective and detoxified bile acid species in bile and feces, along with an overall reduction in hepatotoxic bile acids in liver and serum.

Building on this overview, individual bile acid profiling showed elevated levels of cytoprotective species including TUDCA, GUDCA, and THDCA in bile (**Figure 5F-H, Suppl Figure 5G**). Rodent-specific muricholic acids (αMCA, βMCA, ωMCA, TaMCA, GbMCA) were also elevated, these species exert FXR-antagonistic but hepatoprotective effects (**Figure 5F**). In addition, we observed increases in 7-keto-DCA, a detoxified derivative of DCA, and secondary bile acids (GDCA, TDCA, TLCA, LCA), indicating enhanced microbial conversion and detoxification (**Suppl Figure 5H**).

In feces, despite unchanged total levels (**Figure 4M**), composition also shifted toward detoxified species —including HDCA, HCA, and UDCA (**Figure 5F, I**). TUDCA trended upward but did not reach significance (**Figure 5I**). In contrast, harmful species like DCA and 12-keto-LCA were significantly reduced (**Figure 5J**), suggesting a more tolerable and anti-inflammatory bile acid excretion profile.

In liver and serum, most bile acids—particularly hydrophobic and hepatotoxic ones like TCA, GCA, and CA—were significantly reduced (**Suppl Figure 5I, 5K**), while protective species (TUDCA, CDCA) remained unchanged or slightly decreased without significance (**Suppl Figure 5J, 5L**). This suggests enhanced hepatic export and reduced retention of toxic bile acids without loss of beneficial pools.

The beneficial bile acid handling required coordinated upregulation of export machinery. Analysis of transporter gene expression revealed upregulation of apical bile acid transporters including Abcb11 (BSEP), Abcg5, and Abcg8, promoting canalicular export (**Figure 5K-L**). Basolateral transporters that move bile acids to blood (Slc51a/b) remained unchanged (**Figure 5K-L**), creating a selective“hepatic unloading” pattern that prevents intrahepatic bile acid accumulation. This transport pattern was also seen in the GAN model, where *Slc10a1*(NTCP, facilitating bile acid uptake from blood) and *Abcb11* (BSEP, hepatocyte to bile) (**Figure 5M-N**) were both elevated, supporting efficient bile acid cycling. Lipid transporters (Abca1, Abcg5/8) were also induced (**Figure 5M-N**), facilitating coordinated cholesterol and phospholipid clearance.

Collectively, ZNRF3/RNF43 deletion promotes β-catenin–driven cholesterol rerouting into bile acid synthesis via the alternative pathway, enhances hepatobiliary export (**Figure 5O**), and establishes a detoxified, hepatoprotective bile acid profile. These changes support hepatic unloading, reduce lipid toxicity, and facilitate resolution of MASH.

### ZNRF3 and RNF43 are therapeutic targets for fatty liver diseases

Given the beneficial effects of ZNRF3/RNF43 deletion on MASH regression without enhancing tumorigenesis and injuries, we next evaluated whether transient knockdown of these genes could offer a viable therapeutic strategy.

Chemically stabilized DsiRNAs targeting *Znrf3* and *Rnf43* were formulated into lipid nanoparticles (LNPs) for systemic delivery. To validate knockdown efficiency *in vivo*, a single dose of LNP-packaged non-targeting control or *Znrf3/Rnf43*-targeting DsiRNAs (10 μg each) was administered intravenously to C57BL/6 mice, and livers were harvested after three days for qPCR analysis (**Figure 6A, Suppl Figure 6A**). Over 50% knockdown was achieved for both genes, confirming effective delivery and silencing (**Figure 6B**). CYP2E1 immunostaining confirmed β-catenin activation, with ∼0.5-fold expansion of CYP2E1+ regions and signal extension to periportal zones (**Suppl Figure 6B-C**).

**Figure 6.**
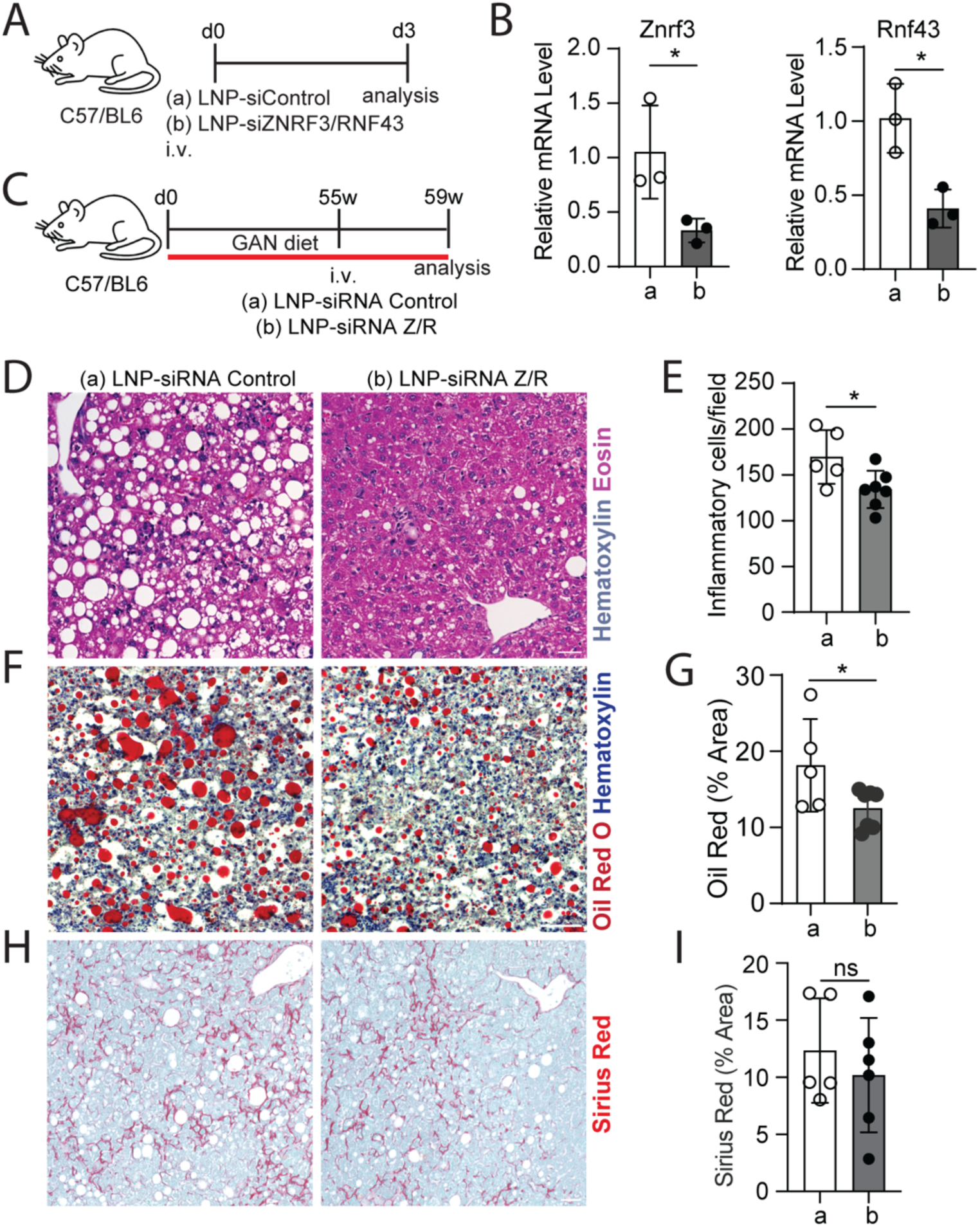
ZNRF3/RNF43 silencing reduces liver lipid accumulation and injury**. (A)** Schematic of LNP-siRNA delivery in WT C57BL/6J mice. **(B)** qPCR analysis of *Znrf3* and *Rnf43* mRNA. **(C)** Schematic of the GAN diet model used to test therapeutic LNP-siZNRF3/RNF43 intervention. **(D–E)** H&E staining and quantification of inflammatory cells. **(F–G)** Oil Red O staining and quantification. **(H–I)** Sirius Red staining and quantification. Data are presented as mean ± s.d., with individual mouse values overlaid as dots. Two-tailed unpaired Student’s t-test. *p < 0.05; ns, not significant. Scale bars: 100 µm (D, F, H).

To evaluate efficacy in established disease, mice were fed the GAN diet for 55 weeks to induce advanced MASH, followed by a single LNP-siRNA injection and continued GAN diet for 4 weeks (**Figure 6C, Suppl Figure 6D**). ZNRF3/RNF43 knockdown led to significant reductions in hepatic steatosis and inflammation (**Figure 6D–G**), with no tumor formation detected. These improvements correlated with localized β-catenin activation (**Suppl Figure 6D–F**). Although fibrosis trended downward, the change did not reach statistical significance **(Figure 6H–I**), indicating that longer treatment duration or improved knockdown efficiency may be necessary. Notably, steatosis and inflammation improvements varied across liver regions, likely reflecting heterogeneous knockdown or limited siRNA persistence. These observations highlight the need for further optimization of siRNA formulation and delivery strategies to maximize therapeutic efficacy.

Collectively, our findings reveal a previously unrecognized therapeutic mechanism whereby transient ZNRF3/RNF43 inhibition activates β-catenin signaling to selectively enhance alternative bile acid synthesis, promoting cholesterol clearance through a detoxified bile acid profile while stimulating hepatic regeneration. This coordinated metabolic and regenerative reprogramming reverses established MASH without safety concerns, establishing ZNRF3/RNF43 as viable therapeutic targets for fatty liver disease treatment.

## Discussion

During homeostasis, a central-to-portal WNT/β-Catenin activity gradient governs liver metabolic zonation, and activation of this pathway is critical for liver regeneration following injury^18,24^. ZNRF3 and RNF43 are E3 ubiquitin ligases that physiologically restrict β-Catenin signaling along this axis to maintain controlled metabolic function^18,24^. In MASLD and MASH, this gradient is disrupted— β-Catenin activity declines globally in hepatocytes while spatial activation expands from the pericentral to periportal zones. Such disruption, also supported by reported data indicating extension of several β-Catenin target genes including GS, and WNT ligands ^22,28^, suggesting that reacquiring of β-Catenin activity at the periportal region coupled with increased detoxification genes could be a potential protective mechanism. Prior studies, including APC knockout mice and Wnt3a rescue experiments in LRP6 ^R611C^ mutant mice, show β-Catenin activation lowers liver lipid content ^29,30^. Our previous data suggested a reduced lipid accumulation upon ZNRF3/RNF43 deletion during homeostasis ^18^. These studies collectively demonstrate that β-Catenin activation at various levels supports hepatic lipid reduction. Here, we found that removing ZNRF3 and RNF43 in hepatocytes during MASLD/MASH reactivates β-Catenin and induces lipid detoxification pathways, enabling in situ lipid catabolism and enhanced regeneration without introducing unexpected pathological burdens.

These effects contrast with observations by Belenguer et al., who reported increased lipid accumulation following ZNRF3/RNF43 deletion in homeostasis and organoid models^19^. Such differences likely stem from divergent experimental contexts—homeostatic versus diseased states, genetic background, analytical time points, or *in vitro* versus *in vivo* settings—especially given the importance of the liver-gut bile acid axis. The controversial data between our previous study and Belenguer et al., also involved in liver regeneration as our prior data showed increased regeneration after partial hepatectomy in ZNRF3/RNF43-deficient mice, whereas Belenguer et al. observed impaired regeneration following chronic CCl_4_ injury^18,19^. Interestingly, in our FAT-MASH model, ZNRF3/RNF43-deficient livers showed intolerance to even low-dose CCl_4_, likely due to global CYP2E1 expression and subsequent generation of toxic metabolites led to hepatocyte death and liver failure (data not shown). This necessitated withdrawal of CCl_4_ post-deletion, further emphasizing that the regenerative response to WNT activation depends on injury context, gene deletion efficiency, and doses/timing CCl_4_ in the model.

WNT/β-Catenin signaling modulates lipid metabolism, inflammation and fibrosis in various contexts. Lipid accumulation contributes significantly to MASLD/MASH pathology by inducing cellular stress, inflammation, and fibrosis, with free cholesterol acting as a key accelerator of disease progression^4,5,31^. Elevated hepatic cholesterol levels are commonly observed in both MASH patients and mouse models^1,32,33^. Redirecting cholesterol into catabolic pathways therefore presents a compelling therapeutic strategy. Although bile acid synthesis is a primary route for cholesterol disposal, impaired bile acid transport in fatty liver disease can lead to accumulation in the liver and serum, exacerbating liver injury^34,35^. Thus, coupling enhanced cholesterol-to-bile acid conversion with improved bile acid export is essential for safely reducing hepatic cholesterol burden in MASH. In our study, four weeks of hepatocyte-specific ZNRF3/RNF43 deletion in established MASH models led to robust regression of steatosis, inflammation, and fibrosis. Our data demonstrate that ZNRF3/RNF43 deletion enhances cholesterol conversion to bile acids via the β-Catenin–dependent alternative pathway (CYP27A1, CYP7B1), without increasing bile acid toxicity. This contrasts with β-Catenin gain-of-function models or APC deletions, which activate both classical (CYP7A1) and alternative (CYP27A1) pathways. Importantly, β-Catenin appears to coordinate bile acid synthesis and export, as shown by our observed upregulation of several key bile acid transporters (NTCP, BSEP), as well as cholesterol and lipid transporters (Abcg5/8) and lipoprotein transporters (Abca1), pointing to an improved bile acid cycling and lipid handling with potential enhanced reverse cholesterol transport (RCT). This stands in contrast to models involving hepatocyte-specific expression of an oncogenic β-Catenin variant, which results in cholestatic liver disease with dysregulated multiple bile acid transporters^16^. In line with our observation, loss of β-Catenin in hepatocytes leads to bile secretory defect, bile canalicular abnormalities and cholestasis, as seen in β-Catenin knockout mice ^36^. Deletion of LGR4/5, key upstream components of canonical WNT signaling, impairs β-Catenin activity and pericentral metabolic zonation, resulting in bile acid accumulation in both the liver and circulation due to compromised bile secretion ^35^. Together, these findings highlight that the physiological outcomes of modulating β-Catenin signaling depend on the nature and extent of activation or inhibition. Such differential regulation likely alters the hepatic transcriptome through interactions with transcription factors such as FXR, LXR, HNF4α, and TCF/LEF1, collectively governing bile acid synthesis, secretion, and transport. Physiological activation of β-Catenin via ZNRF3/RNF43 deletion promotes a targeted and non-pathological bile acid profile, characterized by selective activation of the alternative bile acid synthesis pathway and effective export mechanisms. Our findings highlight the unique therapeutic advantage of modulating β-Catenin through endogenous pathway regulators like ZNRF3/RNF43, enabling metabolic reprogramming without triggering adverse outcomes.

Furthermore, ZNRF3/RNF43 deletion reshapes the bile acid composition across compartments, contributing to a more favorable metabolic profile. Bile acid synthesis and excretion are major catabolic pathways for cholesterol and lipid clearance. Numerus studies have shown that intestinal bile salt hydrolase (BSH) inhibitors, such as riboflavin, caffeic acid phenethyl ester, grape seed extracts, and Theabrownin, enhance lipid metabolism by promoting alternative bile acid synthesis ^6–10^. This effect is particularly associated with reduced hepatic cholesterol and lipid levels and increased expression of CYP7B1, a key enzyme in the alternative pathway. In contrast, CYP8B1 overexpression and the resulting increase12-OH bile acids from the classical pathway have been linked to metabolic disorders such as insulin resistance, type-2 diabetes in both mouse models and human ^37–39^. Notably, reducing 12-OH bile acids through CYP8B1 deficiency elevates the non-12-OH/12-OH bile acid ratio, improving glucose tolerance and protecting against diet-induced obesity, hepatic steatosis, and insulin resistance^40–42^. In our study, ZNRF3/RNF43 deletion led to protective species like TUDCA, GUDCA, and THDCA increased in bile, while hepatotoxic bile acids such as TCA, DCA, and GCA decreased in liver. This pattern of remodeling favors a cytoprotective and anti-inflammatory profile, supporting MASH regression. This detoxification is further complemented by increased expression of lipid transporters (Abca1, Abcg5/8), pointing to enhanced hepatic unloading rather than accumulation.

In addition to its roles in reducing steatosis-based lipid toxicity, alternative bile acid profiling and lipid handling, β-Catenin activation has also been shown to directly suppress hepatic inflammation and fibrosis ^43,44^. Nejak-Bowen et al. demonstrated that β-Catenin can inhibit NF-κB activity by forming a complex with its p65 subunit in hepatocytes; correspondingly, β-Catenin knockout leads to heightened inflammatory responses^44^. Moreover, β-Catenin activation has been shown to attenuate fibrosis in CCl_4_-induced liver injury models, even in the context of relatively modest inflammation responses ^45^. These findings suggest that β-Catenin activation downstream of ZNRF3/RNF43 deletion may contribute to rapid disease regression not only through lipid clearance but also by suppressing fibrotic and inflammatory pathways. Further investigation is warranted to dissect these steatosis-independent mechanisms and their interplay during ZNRF3/RNF43-driven MASH resolution.

The therapeutic potential of these findings is reinforced by our proof-of-concept siRNA experiments. A single dose of LNP-formulated ZNRF3/RNF43 siRNAs reversed steatosis and inflammation in mice on GAN diet for over a year. Importantly, this transient intervention did not lead to tumor formation, suggesting a favorable safety profile. However, to enhance efficacy and durability, further optimization of siRNA properties—including dosage, timing, and in vivo stability—is necessary to achieve more robust and sustained knockdown. Additionally, alternative hepatocyte-specific delivery strategies, such as GalNAc-conjugated siRNAs or inducible AAV-based systems, particularly those allowing fine-tuned stoichiometric targeting of ZNRF3 and RNF43, may further improve therapeutic potency and precision.

Safety is a major concern in regenerative medicine, particularly when targeting highly conserved pathways such as WNT/β-Catenin, which are essential for both development and tissue repair. Unlike constitutive or gain-of-function mutations in β-Catenin that led to sustained and potentially oncogenic signaling, ZNRF3 and RNF43 serve as physiological gatekeepers that dynamically modulate β-Catenin activity during homeostasis and regeneration^18,24^. Prior studies, including our own, have shown that hepatocellular carcinoma (HCC) arises only after prolonged ZNRF3/RNF43 deletion (e.g., one year in homeostasis or seven months with chronic CCl_4_-induced injury) ^18,19^. Moreover, β-Catenin activation alone has not been sufficient to induce HCC in these contexts^20^. This suggests that ZNRF3/RNF43 deletion likely reprograms hepatocyte metabolism and fitness rather than directly transforming cells into tumorigenic states. In both FAT-MASH and GAN diet-induced MASH models, hepatocyte-specific ZNRF3/RNF43 deletion resulted in robust disease regression within four weeks, without evidence of additional liver injury or tumor formation. These outcomes were sustained in a 12-week follow-up, supporting the existence of a safe and effective therapeutic window for transient knockdown. Nonetheless, further investigation is needed to fully assess tumorigenic risk, particularly in models predisposed to HCC and long-term tumor monitoring post treatment.

In summary, our findings demonstrate that transient, physiological reactivation of WNT/β-Catenin signaling via ZNRF3/RNF43 deletion reverses key MASH features, including steatosis, inflammation, and fibrosis. Mechanistically, this therapeutic effect is dependent on β-Catenin and is driven by the selective activation of cholesterol-to-bile acid conversion through the alternative bile acid synthesis pathway. This occurs without triggering oncogenic pathways or hepatotoxicity, distinguishing it from other modes of β-Catenin activation. These insights identify ZNRF3 and RNF43 as promising therapeutic targets for reversing fatty liver disease.

## Limitations of the study

(1) While mouse models of fatty liver disease and MASH demonstrated robust regression following hepatocyte-specific ZNRF3/RNF43 deletion, further studies are needed to assess the relevance and therapeutic potential of targeting these genes in human MASH. (2) The ZNRF3/RNF43–β-Catenin–Cyp27a1 axis appears to facilitate the conversion of cholesterol into bile acids through the alternative pathway, accompanied by enhancements in anti-inflammatory and anti-fibrotic bile acid profiles, bile acid transport, and lipid handling. However, whether ZNRF3/RNF43-mediated β-Catenin activation can reduce fibrosis and inflammation independently of alternative bile acid synthesis remains unclear. (3) Although we observed no tumor formation during the study period, additional long-term safety studies post treatment and HCC-relevant tumorigenesis models are needed to comprehensively evaluate safety across diverse pathological contexts.

## Acknowledgements

We thank Jan Tchorz for providing the ZNRF3/RNF43^fl/fl^ mice. We also thank Scott Friedman, Sarah Millar, Nicole Dubois, Florence Marlow, Michael Rendl, James Bieker, Robert Krauss and Meritxell Huch for their helpful discussions.

## Author contributions

Conceptualization, T.S., F.D.T., S.B.Y. L.Y.; Methodology, F.D.T., S.B.Y. S.R., K.G., A.G., J.C., S.V.K., J.A., H.L., S.S., M.B., E.B., A.L., L.S., D.Z., H.P., F.M., B.G., L.Y, T.S.; Formal analysis, F.D.T., S.B.Y. S.R., K.G., A.G., J.C., S.V.K., J.A., S.S., L.Y, T.S.; Investigation, F.D.T., S.B.Y. S.R., K.G., A.G., J.C., S.V.K., J.A., H.L., S.S., M.B., E.B., A.L., L.S., D.Z., H.P., F.M., L.Y, T.S.; Supervision, A.L., L.Y., B.C., H.Z., Y.D., S.M., T.S.; Writing – original draft, S.B.Y., F.D.T., L.Y., T.S..

## STAR Methods

### Resource availability

#### Lead contact

For further information and requests for resources and reagents, please contact the Lead Contact, Tianliang Sun

#### Materials availability

This study did not produce any new or unique reagents.

#### Data and software availability

Bulk RNA-seq data are available at the NIH Sequence Read Archive once manuscript is published. All software and algorithms use for data analyses are previously published and appropriately referenced. Further details on data analysis reported in this paper are available from the Lead Contact, Tianliang Sun, upon request.

### Experimental model and subject details

#### Mouse models

ZNRF3/RNF43^fl/fl^ mice ^24^ and R26-STOP-EGFP mice ^46^ were described previously. ZNRF3/RNF43/CTNNB1 triple floxed mice ZNRF3/RNF43/CTNNB1^fl/fl^ were generated by crossing ZNRF3/RNF43^fl/fl^ mice with CTNNB1 floxed mice CTNNB1^fl/fl^, a kind gift from Dr. Rendl lab (Icahn School of Medicine at Mount Sinai) (Jax#022775). Hepatocyte specific ZNRF3/RNF43 deletion mice ZNRF3/RNF43^ΔHEP^ and ZNRF3/RNF43/CTNNB1 deletion mice ZNRF3/RNF43/CTNNB1^ΔHEP^ were obtained by AAV8-TBG-Cre injection to ZNRF3/RNF43^fl/fl^ or ZNRF3/RNF43/CTNNB1^fl/fl^ mice respectively. C57BL/6J mice (8–12 weeks old) were obtained from The Jackson Laboratory.

#### In Vivo Mouse Studies

The FAT-MASH model^5,23^ in ZRNF3/RNF43^ΔHEP^ mice was induced by feeding age-matched male ZNRF3/RNF43^fl/fl^ mice with high-fat Western diet (20–23% fat by weight, 40–45% kcal from fat), rich in milk fat (saturated fatty acids >60% of total fat) and 1.25% cholesterol (Envigo, Cat# TD.120528), with glucose/fructose-supplemented drinking water, and chronic weekly administration of a low-dose carbon tetrachloride (CCl₄; 0.02 mL/kg body weight) for 20 weeks followed by a single dose of 5×10E10 genome copies (GC) of AAV8-TBG-EGFP (control) (Addgene Cat# 105535-AAV8) or AAV8-TBG-Cre (ZRNF3/RNF43^ΔHEP^) (Addgene #107787-AAV8)injection i.v. per mouse, and the mice were continued on the same diet and water but withdraw CCl4 treatment was withdrawn for additional 4 weeks. The GAN diet (contains 40% kcal from fat (primarily palm oil), 20% kcal from fructose, and 2% cholesterol) (Research Diets Inc., Cat# D09100310) induced MASH model in Rosa26-LSL-EGFP, ZRNF3/RNF43^ΔHEP^ or ZNRF3/RNF43/CTNNB1^ΔHEP^ mice was induced by feeding with age-matched adult males and females for 16, 28 or 35 weeks followed by a dose of 5×10E10 GC of AAV-TBG-EGFP (control) or AAV8-TBG-Cre (ZRNF3/RNF43^ΔHEP^) injection i.v. per mouse, and maintained the mice on the same diet for an additional 2-12 weeks before analysis. For the LNP-siRNA knockdown efficiency study, 8-12 weeks old C57/BL6 male mice were injected i.v. 200 ul of LNP-siRNA containing either a negative control (20ug) or equimolar amounts of si-ZNRF3 and si-RNF43 (10ug each) followed by liver sampling and analysis after 3 days. For the LNP-siRNA study in the GAN diet-induced MASH mode, age-match adult (8-12 weeks) C57/BL6 mice were fed in GAN diet for 55 weeks followed by injection 200 μl of LNP-siRNA containing either a negative control (20 μg) or equimolar amounts of si-ZNRF3 and si-RNF43 (10 μg each) followed by additional 4 weeks on the same diet before the sampling. In experiments involving GAN diet, body weight (BW) was measured weekly for each mouse and food intake was assessed weekly on a per-cage basis by weighing the food reservoir at the start and end of each week. Weekly food intake was normalized to the number of mice per cage and body weight gain was applied to assess body weight change through the study.

Mice were permanently identified at weaning (postnatal day 21) using the GEMS (Genetically Engineered Mouse Subcommittee) ear notching system, encoding ID numbers using predefined left/right ear notch positions. Age-matched adult male and female mice were randomly assigned to experimental groups in balanced numbers. Animals were housed in the Icahn Building Mouse Facility under a 12-hour light/dark cycle in cages with nesting enrichment. Breeding cages were limited to two mice and experimental cages to five. Single housing was used only when necessary (e.g., post-operative recovery or aggression). The facility adheres to institutional biosecurity and pathogen exclusion protocols, separating breeding, procedural, and surgical areas. All procedures were approved by the Institutional Animal Care and Use Committee (IACUC) at the Icahn School of Medicine at Mount Sinai. Mice were monitored regularly for health and had *ad libitum* access to food and water.

## Method details

### Histological staining and imaging

Tissue samples were freshly frozen in O.C.T. embedding medium (Tissue TEK) or fixed for 28 hours in 10% buffered formalin and embedded in paraffin. Primary antibodies used in this study were Rabbit anti-CYP2E1 (Sigma-Aldrich, Cat# HPA009128), Rabbit anti-Ki67 (Abcam, Cat# ab15580), Rat anti-Ki-67 (eBioscience, Cat# 14-5698-82), Mouse anti-E-Cadherin (BD Biosciences, Cat# 610181), Chicken anti-Albumin (Sigma-Aldrich, Cat# SAB3500217), Rabbit anti-Cleaved Caspase-3 (Cell Signaling Technology, Cat# 9661S; RRID: AB_2341188), Rat anti-CK19 (Developmental Studies Hybridoma Bank, Cat# TROMA-III; RRID: AB_2133570), Goat anti-GFP (Abcam, Cat# ab6673; RRID: AB_305643), Rabbit anti-CYP1A2 (Abcam, Cat# ab22717), and Rabbit anti-CYP27A1 (Abcam, Cat# ab126785). Alexa DyLight-or Cy-coupled (Jackson ImmunoResearch) secondary antibodies were used for primary antibody detection, and either Hematoxylin Gill III (Sigma-Aldrich, Cat# 65067-75) or 4’-6-diamidino-2-phenylindole (DAPI; Sigma-Aldrich, Cat# D9542) were used for counterstaining. For non-fluorescent stainings, Histofine Simple Stain MAX PO secondary reagents Anti-rabbit (Nichirei Biosciences Inc., Cat# 414341F) and Anti-rat (Nichirei Biosciences Inc., Cat# 414311F) and ImmPACT DAB substrate kit (Vector Laboratories, Cat# SK-4105) were used. For Oil Red O staining, slides from fresh frozen samples were fixed in formalin for 10 minutes, rinsed under tap water for 2minutes and dipped in 60% 2-propanol (Sigma-Aldrich, Cat# I9516) for 5 minutes. Slides were stained in the Oil Red O working solution for 10 minutes, rinsed in MilliQ water until the background cleared, and counterstained with Hematoxylin Gill III (Sigma-Aldrich, Cat# 65067-75) for 1 minute followed by sealing with glycerol (Sigma-Aldrich, Cat# G5516). For Sirius Red staining, formalin-fixed paraffin-embedded (FFPE) sections were deparaffinized and stained in a 0.01% Fast Green (Sigma-Aldrich, Cat# F7258) solution at RT for 1h and in 0.04% Fast Green plus 0.1% Direct Red 80 (Sigma-Aldrich, Cat# 365548-5G) solution for 1 h on a shaker. Slides were washed with distilled water, dehydrolyzed and mounted using Permount (ThermoFisher, Cat# SP15-100).

For representative images, whole-liver lobes were examined histologically in multiple replicates. Image analysis and quantification of Oil Red O, Sirius Red staining on cross sections of liver lobes was performed using the Color Deconvolution plugin from the Fiji ImageJ software. Data was presented as percent positive stained area per total analysis area or percent positive cells per high power field image. FISH for human AXIN2 probe (ACD, Cat# 400331-C3) were performed accordingly to the manufacturer’s instruction. Briefly, 5 µm FFPE human liver sections were deparaffinized, quenched before the antigen retrieval by boiling for 20minutes at 98 °C in a microwave host processor. Slides were dried for protease digestion with RNAscope Protease Plus (ACD, part of Cat# 323100) at 40°C for 30 minutes before incubating with AXIN2-C2 probe for 2 hours at 40 °C. Signal amplification was achieved using the RNAscope Multiplex Fluorescent Reagent Kit v2 (ACD, Cat# 323100). Signal detection was performed using Opal 570 (Akoya; diluted 1:1000 in TSA buffer; ACD, Cat# 322810). CK19 and GS co-staining was performed overnight followed by secondary antibody and DAPI incubation and mounted by FluoroSave mounting medium (Millipore, Cat# 345789). The percent FISH signal per analyses area was quantified using Fiji ImageJ software.

Brightfield, immunofluorescence, and RNA fluorescence in situ hybridization (FISH) images were acquired using a Stand Axio Observer 7 Microscope (Carl Zeiss AG, 431007-9904-000) equipped with a motorized stage, high-resolution CMOS camera, and an LED-based fluorescence illumination system. Imaging was performed using 10X and 20X objectives, selected based on the experimental objectives. Brightfield images were acquired using standardized exposure and white balance settings. Fluorescence imaging utilized filter sets optimized for Cy2 (green), Cy3 (red), and Cy5 (far-red) fluorophores, selected based on the specific emission spectra of each fluorophore.

### Transmission Electron Microscopy (TEM) Sample Preparation

Liver tissues were prepared for transmission electron microscopy (TEM) using established fixation and embedding protocols. Tissue blocks containing regions of interest were dissected and rinsed in 0.1 M sodium cacodylate buffer (Electron Microscopy Sciences, Cat# 11652), followed by post-fixation with 1% osmium tetroxide (Electron Microscopy Sciences, Cat# 19100) in the same buffer. Samples were dehydrated in graded ethanol (70%–100%), transitioned into propylene oxide (Electron Microscopy Sciences, Cat# 20401), and infiltrated with Epon resin (Electron Microscopy Sciences, Cat# 14120). Resin blocks were polymerized in a vacuum oven at 60°C for 48 hours. Semithin sections (1 μm) were stained with toluidine blue (Sigma-Aldrich, Cat# T3260) to localize regions of interest. Ultrathin sections (80 nm) were cut with a diamond knife using a Leica UCT ultramicrotome (Leica Microsystems) and mounted on copper grids with Coat-Quick Glo adhesive (Electron Microscopy Sciences, Cat# 10310). Grids were counterstained with 2% uranyl acetate (Electron Microscopy Sciences, Cat# 22400) and lead citrate (Electron Microscopy Sciences, Cat# 17800). TEM samples were imaged with a Hitachi 7700 transmission electron microscope (Hitachi High-Technologies Corp.) equipped with an Advantage CCD camera (Advanced Microscopy Techniques). Image brightness and contrast were adjusted using Adobe Photoshop 2022 (Adobe Systems, RRID: SCR_014199).

### Bulk RNA-seq of MASH Livers and Count Generation

Total RNA was extracted from snap-frozen liver tissue of ZNRF3/RNF43^fl/fl^ (n = 3) and ZNRF3/RNF43^ΔHep^ (n = 2) mice using the RNeasy Plus Mini Kit (QIAGEN, Cat# 74134), following the manufacturer’s protocol. RNA concentration and integrity were assessed using the Agilent RNA 6000 Nano Kit (Agilent Technologies, Cat# 5067-1511). Bulk RNA sequencing was performed by Novogene Corporation Inc. (Sacramento, CA, US). Messenger RNA was purified from total RNA using poly-T oligo-attached magnetic beads. Library construction was performed using the Illumina TruSeq RNA Sample Prep Kit v2 (Illumina, Cat# RS-122-2001). Libraries were quantified using Qubit fluorometry and real-time PCR, with fragment size distribution confirmed by Bioanalyzer. Sequencing was performed on an Illumina NovaSeq 6000 platform, yielding paired-end 150 bp reads. Each sample yielded between 44 and 52 million clean reads, with a GC content of between 49% and 50%. The average sequencing error rate was 0.03%, with >97% of bases achieving Q20 and >92% achieving Q30 quality scores. Alignment to the *Mus musculus* mm39 reference genome was performed using STAR v2.7.11a, resulting in high mapping efficiency with total mapping rates of 92.0–94.6% and unique mapping rates of 81.2–85.8%. Properly paired reads ranged from 77.8% to 82.5%. The read distribution analysis revealed that most reads aligned to exonic regions, with smaller proportions mapping to intronic or intergenic regions. Transcript quantification was conducted using an in-house pipeline based on Ensembl gene annotations (release 98), as previously described ^47^. Gene-level raw counts were generated using Ensembl gene IDs. Differential expression analysis was performed using DESeq2 v1.42.0 in R v4.4.1, following the normalization of raw counts.

### snRNA-Seq data processing and correlation between hepatocyte gene expression and clinical phenotypes

For downstream analysis of the human liver single-nucleus RNA-seq dataset (GSE202379; n = 47 liver samples), we utilized the preprocessed Seurat object provided by Gribben et al. (GSE202379_SeuratObject_AllCells.rds.gz). The dataset includes samples spanning healthy control (n = 4), MASLD (n = 7), MASH without cirrhosis (n = 27), MASH with cirrhosis (n = 4), and end-stage liver disease (n = 5)^22^. The dataset was previously quality filtered, normalized using the SCTransform method, integrated across samples with the Harmony algorithm (version 1.2.3), and annotated into major hepatic and non-parenchymal cell types, including hepatocytes, cholangiocytes, endothelial cells, macrophages, hepatic stellate cells, lymphocytes, neutrophils, and B cells. To investigate the relationship between hepatocyte-specific gene expression and clinical disease parameters, the average expression of selected genes per patient was computed. For each gene and clinical variable, we summarized patient-level expression and compared distributions across clinical categories, including Disease Status, Steatosis Grade, Ballooning, Inflammation, and Fibrosis Stage. Visualizations were generated using boxplots overlaid with individual data points, color-coded by disease status. All statistical analyses and visualizations were performed in R (version 4.4.1) using Seurat (version 5.1.0) and ggplot2 (version 3.5.2). All processing steps followed the methodology described in the original publication. Additional analytical details and custom code used for subsetting, visualization, and differential expression analysis are available upon request.

### Histopathological scoring of H&E slides

Histopathological analysis from H&E staining was performed by a blinded pathologist according to the rodent MASLD scoring system^25^. Briefly, the Steatosis Score (SS) was calculated for each mouse as the sum of the scores of macrovesicular steatosis, microvesicular steatosis, and hepatocellular hypertrophy, scored as 0 (<5% of the microscopic field), 1 (5–33%), 2 (33–66%), or 3 (>66%). Lobular inflammation was classified as displaying normal (0, <0.5 inflammatory foci/field), slight (1, 0.5–1.0 foci/field), moderate (2, 1.0–2.0 foci/field), or severe (3, >2.0 foci/field) inflammation.

### Formulation and Preparation of siRNA-Loaded Lipid Nanoparticles (LNPs)

N1, N3, N5-tris(3-(didodecylamino) propyl) benzene-1,3,5-tri carboxamide (TT3) was kindly provided by the Dong Research Group. Additional lipid components, including cholesterol (Avanti Polar Lipids, Cat# 700000P), 1,2-dioleoyl-sn-glycero-3-phosphoethanolamine (DOPE; Avanti Polar Lipids, Cat# 850725P), and 1,2-dimyristoyl-rac-glycero-3-methoxypolyethylene glycol-2000 (DMG-PEG2k; Avanti Polar Lipids, Cat# 880151P), were purchased from Avanti Polar Lipids (Alabaster, AL, USA). siRNA-loaded lipid nanoparticles (LNPs) were formulated using the Rapid Nanomedicine System INano L+ (Micro&Nano Biologics Technology Ltd.), following a previously described protocol^48,49^. Briefly, lipids were dissolved in absolute ethanol at a molar ratio of TT3/DOPE/cholesterol/DMG-PEG2k = 20:30:40:0.75 and rapidly mixed with an aqueous solution of siRNA (targeting ZNRF3 and RNF43, sequence details in Key Resources Table) using microfluidic mixing, maintaining a lipid-to-siRNA mass ratio of 10:1. The resulting nanoparticles were dialyzed against phosphate-buffered saline (PBS, pH 7.4, ThermoFisher Scientific, Cat# 10010-023) for 80 minutes at room temperature using Slide-A-Lyzer Dialysis Cassettes (ThermoFisher, Cat# 66380; 10K MWCO). LNPs had a mean hydrodynamic diameter of ∼80 nm and were characterized for size and homogeneity by dynamic light scattering (DLS) before intravenous injection into C57BL/6 mice. All steps were carried out under RNase-free and sterile conditions. Final LNP formulations were stored on ice and administered within 24 hours of preparation.

### Liver Lipids Measurement

Liver lipids were isolated using the Bligh and Dyer method^50^. Briefly, frozen liver tissues (50 to 100 mg) were homogenized in 0.5 ml PBS. Liver homogenates (200 µl) were mixed with 6 ml of chloroform: methanol: PBS (8:4:3) in 20 ml scintillation vial by vertexing. The mixture was centrifuged at 2,000 x g for 10 min to separate into different layers. The organic bottom layer was carefully transferred to a 20 ml scintillation vial using a glass Pasteur pipet, avoiding contamination from other layers. The leftover layers were mixed with 5 ml chloroform: methanol: PBS (86:14:1) to separate the organic layer once more. The organic layers were combined and carefully blown down under nitrogen gas. The lipids were solubilized with 1 ml of 5% Triton X-100 in chloroform and blown down again with nitrogen gas. The lipids were then re-solubilized in 3 ml of ddH2O and subjected to total, free cholesterol, and TG analysis.

### Immunoblotting

Liver tissues were snap-frozen and homogenized in RIPA lysis buffer supplemented with protease inhibitors before concentrations were measured using Pierce BCA Protein Assay Kit(ThermoFisher, Cat# 23225). Protein was loaded onto NuPAGE™ 4–12% Bis-Tris gels (ThermoFisher, Cat# NP0323BOX) for electrophoretic separation, PVDF membrane transfer, blocking, then incubated with the following primary antibodies: Anti-CYP27A1 (Rabbit polyclonal, Abcam, Cat# ab126785, 1:2000); Anti-CYP7B1 (Rabbit Polyclonal Antibody, Proteinbiotech, 1:1000), Anti-CYP1A2 (Rabbit polyclonal, Abcam, Cat# ab22717, 1:1000); Anti-GAPDH (Mouse monoclonal, Proteintech, Cat# 60004-1-Ig, 1:50000). HRP-conjugated secondary antibodies: Anti-rabbit IgG-HRP (Cell Signaling Technology, Cat# 7074, 1:2000); Anti-mouse IgG-HRP (Bio-Rad, Cat# 1706516, 1:3000) were used. Signal detection was performed using enhanced chemiluminescence, and images were acquired with the Bio-Rad ChemiDoc™ Imaging System.

### RNA extraction, reverse transcription and quantitative RT-PCR (qPCR)

Total RNA was extracted from snap-frozen liver tissue using the RNeasy Plus Mini Kit (QIAGEN, Cat# 74134). Reverse transcription was performed using the High-Capacity cDNA Reverse Transcription Kit (ThermoFisher, Cat# 4368813). Quantitative PCR was performed using iTaq™ Universal SYBR® Green Supermix (Bio-Rad, Cat# 1725121) and gene-specific primers in technical triplicates. PCR reactions were run on a Roche light cycler 480 Real-Time PCR detection system and data processing was conducted using the ΔΔCt method and normalized to β-actin.

### Quantitative Measurement of Serum Alanine Aminotransferase (ALT) activity

ALT activity was measured using the ALT (GPT) Reagent Set (Teco Diagnostics, Cat# A524-150) by adapting the manufacturer’s protocol for 96-well plates. ALT activity was calculated using a manufacturer-defined conversion factor derived from the NADH extinction coefficient. Quality control was performed using standard and abnormal control serum; samples with visible hemolysis or values outside the acceptable control range were excluded.

### Quantitative measurement of Cholesterol and Triglycerides level in serum and feces

Total serum cholesterol was determined using the Cholesterol Reagent Kit (Teco Diagnostics, Cat# C518-120) according to the adapted 96-well plate protocol. Absorbance was measured at 525 nm, and cholesterol concentrations were calculated using a standard curve generated from known cholesterol standards.

Serum triglyceride levels were quantified using the Triglyceride GPO Liquid Reagent Kit (Teco Diagnostics, Cat# T7532-150) according to the microplate protocol. Triglyceride concentrations were interpolated from a standard curve prepared from known standards. Assay performance was validated using normal and abnormal reference sera. Samples showing hemolysis, lipemia, icterus, or glycerol contamination were excluded from analysis.

Lipids from feces were extracted from frozen stool samples using a biphasic chloroform– methanol-water system. Quantitative determination of cholesterol and triglycerides was conducted using Teco Diagnostics colorimetric kits, as described in the respective sections.

### Quantitative Analysis of Bile Acids in Liver, Serum, Bile, and Feces by LC-MS/MS

Bile acid concentrations in liver, serum, bile, and fecal samples were quantified using a standardized liquid chromatography-tandem mass spectrometry (LC-MS/MS) protocol. Liver tissue (1 mg) was homogenized in 150 µL of acetonitrile:methanol (1:1, v/v; Sigma-Aldrich, Acetonitrile Cat# 34967, Methanol Cat# 179337-4L). Twenty microliters of homogenate were combined with 30 µL of acetonitrile:methanol (1:1, v/v), 50 µL of LC-MS grade water (Sigma-Aldrich, Cat# 270733), and 20 µL of internal standard (ISTD) mix, followed by centrifugation at 16,000g for 5 minutes. Serum and bile samples (20 µL) were first precipitated with 200 µL of acetonitrile:methanol (1:1, v/v), and then 20 µL of the resulting supernatant was mixed with 30 µL of acetonitrile:methanol, 50 µL of water, and 20 µL of ISTD, vortexed, and centrifuged at 16,000g for 5 minutes. For fecal samples, 1 mg of dried pellet was homogenized in 550 µL of acetonitrile:methanol (1:1, v/v; Sigma-Aldrich) using a bead-based disruption system. Twenty microliters of the extract were then processed as described above. In all cases, 60 μL of the final supernatant was transferred into glass inserts in 2 mL autosampler vials (ThermoFisher, Cat# C4010-630) and capped for analysis. Internal standards consisted of 500 nM of deuterated bile acids—including d4-CA, d4-CDCA, d4-DCA, d4-LCA, d4-GCA, d4-GDCA, d4-GLCA, d4-TCA, d4-TCDCA, d4-TLCA, d4-CA-3-S, d4-LCA-3-S, and d4-CDCA-3-S—prepared in methanol:acetonitrile:water (20:30:50, v/v/v; Sigma-Aldrich). Calibration curves with nine concentration points (0.3, 1, 3, 10, 30, 100, 300, 1000, 3000 nM) were generated for each batch.

LC-MS/MS was performed on a Shimadzu LCMS-8060 CL triple quadrupole mass spectrometer (Shimadzu, Japan) equipped with an electrospray ionization (ESI) source. Chromatographic separation was performed using a Hypersil GOLD aQ C18 column (100 × 2.1 mm, 1.9 µm; ThermoFisher, Cat# 25302-102130). The LC system consisted of a CBM-20A CL communications module, two LC-30AD CL pumps, DGU-20A3R and DGU-20A5A degassers, a SIL-30AC CL autosampler, and a CTO-20A CL column oven (Shimadzu, part of the LCMS-8060 CL platform). Data acquisition and quantification were performed using LabSolutions Insight software (Shimadzu, Japan).

## Quantification and Statistical Analysis

All data are presented as mean ± standard deviation (s.d.). n refers to biological replicates. Statistical analyses were conducted using GraphPad Prism software (GraphPad Software; RRID:SCR_002798). No statistical method was used to predetermine sample size, and no animals or data points were excluded from analysis. Experimental groups were not randomized or blinded. Data are expected to follow a normal distribution. For comparisons between two groups, two-tailed unpaired Student’s t tests were applied as appropriate. For analyses involving more than two groups, one-way ANOVA was performed, followed by Sidak’s multiple comparisons test. Complete statistical details, including test types, degrees of freedom, and exact p-values, are provided in the corresponding figure legends.

## References

1. Horn, C.L., Morales, A.L., Savard, C., Farrell, G.C., and Ioannou, G.N. (2022). Role of Cholesterol-Associated Steatohepatitis in the Development of NASH. Hepatol Commun 6, 12–35. 10.1002/hep4.1801.

2. Van Rooyen, D.M., Larter, C.Z., Haigh, W.G., Yeh, M.M., Ioannou, G., Kuver, R., Lee, S.P., Teoh, N.C., and Farrell, G.C. (2011). Hepatic free cholesterol accumulates in obese, diabetic mice and causes nonalcoholic steatohepatitis. Gastroenterology 141, 1393–1403, 1403 e1391-1395. 10.1053/j.gastro.2011.06.040.

3. Rom, O., Liu, Y., Liu, Z., Zhao, Y., Wu, J., Ghrayeb, A., Villacorta, L., Fan, Y., Chang, L., Wang, L., et al. (2020). Glycine-based treatment ameliorates NAFLD by modulating fatty acid oxidation, glutathione synthesis, and the gut microbiome. Sci Transl Med 12. 10.1126/scitranslmed.aaz2841.

4. Das, S., Finney, A.C., Anand, S.K., Rohilla, S., Liu, Y., Pandey, N., Ghrayeb, A., Kumar, D., Nunez, K., Liu, Z., et al. (2024). Inhibition of hepatic oxalate overproduction ameliorates metabolic dysfunction-associated steatohepatitis. Nat Metab 6, 1939–1962. 10.1038/s42255-024-01134-4.

5. Tsuchida, T., Lee, Y.A., Fujiwara, N., Ybanez, M., Allen, B., Martins, S., Fiel, M.I., Goossens, N., Chou, H.I., Hoshida, Y., and Friedman, S.L. (2018). A simple diet-and chemical-induced murine NASH model with rapid progression of steatohepatitis, fibrosis and liver cancer. J Hepatol 69, 385–395. 10.1016/j.jhep.2018.03.011.

6. Huang, F., Zheng, X., Ma, X., Jiang, R., Zhou, W., Zhou, S., Zhang, Y., Lei, S., Wang, S., Kuang, J., et al. (2019). Theabrownin from Pu-erh tea attenuates hypercholesterolemia via modulation of gut microbiota and bile acid metabolism. Nat Commun 10, 4971. 10.1038/s41467-019-12896-x.

7. Fang, S., Suh, J.M., Reilly, S.M., Yu, E., Osborn, O., Lackey, D., Yoshihara, E., Perino, A., Jacinto, S., Lukasheva, Y., et al. (2015). Intestinal FXR agonism promotes adipose tissue browning and reduces obesity and insulin resistance. Nat Med 21, 159–165. 10.1038/nm.3760.

8. Smith, K., Zeng, X., and Lin, J. (2014). Discovery of bile salt hydrolase inhibitors using an efficient high-throughput screening system. PLoS One 9, e85344. 10.1371/journal.pone.0085344.

9. Dong, Z., and Lee, B.H. (2018). Bile salt hydrolases: Structure and function, substrate preference, and inhibitor development. Protein Sci 27, 1742–1754. 10.1002/pro.3484.

10. Seo, K.H., Bartley, G.E., Tam, C., Kim, H.S., Kim, D.H., Chon, J.W., Kim, H., and Yokoyama, W. (2016). Chardonnay Grape Seed Flour Ameliorates Hepatic Steatosis and Insulin Resistance via Altered Hepatic Gene Expression for Oxidative Stress, Inflammation, and Lipid and Ceramide Synthesis in Diet-Induced Obese Mice. PLoS One 11, e0167680. 10.1371/journal.pone.0167680.

11. Apte, U., Singh, S., Zeng, G., Cieply, B., Virji, M.A., Wu, T., and Monga, S.P. (2009). Beta-catenin activation promotes liver regeneration after acetaminophen-induced injury. Am J Pathol 175, 1056–1065. 10.2353/ajpath.2009.080976.

12. Planas-Paz, L., Sun, T., Pikiolek, M., Cochran, N.R., Bergling, S., Orsini, V., Yang, Z., Sigoillot, F., Jetzer, J., Syed, M., et al. (2019). YAP, but Not RSPO-LGR4/5, Signaling in Biliary Epithelial Cells Promotes a Ductular Reaction in Response to Liver Injury. Cell Stem Cell 25, 39–53 e10. 10.1016/j.stem.2019.04.005.

13. Sun, T., Pikiolek, M., Orsini, V., Bergling, S., Holwerda, S., Morelli, L., Hoppe, P.S., Planas-Paz, L., Yang, Y., Ruffner, H., et al. (2020). AXIN2(+) Pericentral Hepatocytes Have Limited Contributions to Liver Homeostasis and Regeneration. Cell Stem Cell 26, 97–107 e106. 10.1016/j.stem.2019.10.011.

14. Colnot, S., Decaens, T., Niwa-Kawakita, M., Godard, C., Hamard, G., Kahn, A., Giovannini, M., and Perret, C. (2004). Liver-targeted disruption of Apc in mice activates beta-catenin signaling and leads to hepatocellular carcinomas. Proc Natl Acad Sci U S A 101, 17216–17221. 10.1073/pnas.0404761101.

15. Gougelet, A., Torre, C., Veber, P., Sartor, C., Bachelot, L., Denechaud, P.D., Godard, C., Moldes, M., Burnol, A.F., Dubuquoy, C., et al. (2014). T-cell factor 4 and beta-catenin chromatin occupancies pattern zonal liver metabolism in mice. Hepatology 59, 2344–2357. 10.1002/hep.26924.

16. Lemberger, U.J., Fuchs, C.D., Karer, M., Haas, S., Stojakovic, T., Schofer, C., Marschall, H.U., Wrba, F., Taketo, M.M., Egger, G., et al. (2016). Hepatocyte specific expression of an oncogenic variant of beta-catenin results in cholestatic liver disease. Oncotarget 7, 86985–86998. 10.18632/oncotarget.13521.

17. Lemberger, U.J., Fuchs, C.D., Schofer, C., Bileck, A., Gerner, C., Stojakovic, T., Taketo, M.M., Trauner, M., Egger, G., and Osterreicher, C.H. (2018). Hepatocyte specific expression of an oncogenic variant of beta-catenin results in lethal metabolic dysfunction in mice. Oncotarget 9, 11243–11257. 10.18632/oncotarget.24346.

18. Sun, T., Annunziato, S., Bergling, S., Sheng, C., Orsini, V., Forcella, P., Pikiolek, M., Kancherla, V., Holwerda, S., Imanci, D., et al. (2021). ZNRF3 and RNF43 cooperate to safeguard metabolic liver zonation and hepatocyte proliferation. Cell Stem Cell 28, 1822–1837 e1810. 10.1016/j.stem.2021.05.013.

19. Belenguer, G., Mastrogiovanni, G., Pacini, C., Hall, Z., Dowbaj, A.M., Arnes-Benito, R., Sljukic, A., Prior, N., Kakava, S., Bradshaw, C.R., et al. (2022). RNF43/ZNRF3 loss predisposes to hepatocellular-carcinoma by impairing liver regeneration and altering the liver lipid metabolic ground-state. Nat Commun 13, 334. 10.1038/s41467-021-27923-z.

20. Harada, N., Miyoshi, H., Murai, N., Oshima, H., Tamai, Y., Oshima, M., and Taketo, M.M. (2002). Lack of tumorigenesis in the mouse liver after adenovirus-mediated expression of a dominant stable mutant of beta-catenin. Cancer Res 62, 1971–1977.

21. Zhu, C., Tabas, I., Schwabe, R.F., and Pajvani, U.B. (2021). Maladaptive regeneration - the reawakening of developmental pathways in NASH and fibrosis. Nat Rev Gastroenterol Hepatol 18, 131–142. 10.1038/s41575-020-00365-6.

22. Gribben, C., Galanakis, V., Calderwood, A., Williams, E.C., Chazarra-Gil, R., Larraz, M., Frau, C., Puengel, T., Guillot, A., Rouhani, F.J., et al. (2024). Acquisition of epithelial plasticity in human chronic liver disease. Nature 630, 166–173. 10.1038/s41586-024-07465-2.

23. Vacca, M., Kamzolas, I., Harder, L.M., Oakley, F., Trautwein, C., Hatting, M., Ross, T., Bernardo, B., Oldenburger, A., Hjuler, S.T., et al. (2024). An unbiased ranking of murine dietary models based on their proximity to human metabolic dysfunction-associated steatotic liver disease (MASLD). Nat Metab 6, 1178–1196. 10.1038/s42255-024-01043-6.

24. Planas-Paz, L., Orsini, V., Boulter, L., Calabrese, D., Pikiolek, M., Nigsch, F., Xie, Y., Roma, G., Donovan, A., Marti, P., et al. (2016). The RSPO-LGR4/5-ZNRF3/RNF43 module controls liver zonation and size. Nat Cell Biol 18, 467–479. 10.1038/ncb3337.

25. Liang, W., Menke, A.L., Driessen, A., Koek, G.H., Lindeman, J.H., Stoop, R., Havekes, L.M., Kleemann, R., and van den Hoek, A.M. (2014). Establishment of a general NAFLD scoring system for rodent models and comparison to human liver pathology. PLoS One 9, e115922. 10.1371/journal.pone.0115922.

26. Hao, H.X., Xie, Y., Zhang, Y., Charlat, O., Oster, E., Avello, M., Lei, H., Mickanin, C., Liu, D., Ruffner, H., et al. (2012). ZNRF3 promotes Wnt receptor turnover in an R-spondin-sensitive manner. Nature 485, 195–200. 10.1038/nature11019.

27. Koo, B.K., Spit, M., Jordens, I., Low, T.Y., Stange, D.E., van de Wetering, M., van Es, J.H., Mohammed, S., Heck, A.J., Maurice, M.M., and Clevers, H. (2012). Tumour suppressor RNF43 is a stem-cell E3 ligase that induces endocytosis of Wnt receptors. Nature 488, 665–669. 10.1038/nature11308.

28. Zhou, Y., Zhao, Y., Carbonaro, M., Chen, H., Germino, M., Adler, C., Ni, M., Zhu, Y.O., Kim, S.Y., Altarejos, J., et al. (2024). Perturbed liver gene zonation in a mouse model of non-alcoholic steatohepatitis. Metabolism 154, 155830. 10.1016/j.metabol.2024.155830.

29. Go, G.W., Srivastava, R., Hernandez-Ono, A., Gang, G., Smith, S.B., Booth, C.J., Ginsberg, H.N., and Mani, A. (2014). The combined hyperlipidemia caused by impaired Wnt-LRP6 signaling is reversed by Wnt3a rescue. Cell Metab 19, 209–220. 10.1016/j.cmet.2013.11.023.

30. Wang, S., Song, K., Srivastava, R., Dong, C., Go, G.W., Li, N., Iwakiri, Y., and Mani, A. (2015). Nonalcoholic fatty liver disease induced by noncanonical Wnt and its rescue by Wnt3a. FASEB J 29, 3436–3445. 10.1096/fj.15-271171.

31. Clapper, J.R., Hendricks, M.D., Gu, G., Wittmer, C., Dolman, C.S., Herich, J., Athanacio, J., Villescaz, C., Ghosh, S.S., Heilig, J.S., et al. (2013). Diet-induced mouse model of fatty liver disease and nonalcoholic steatohepatitis reflecting clinical disease progression and methods of assessment. Am J Physiol Gastrointest Liver Physiol 305, G483–495. 10.1152/ajpgi.00079.2013.

32. Ioannou, G.N. (2016). The Role of Cholesterol in the Pathogenesis of NASH. Trends Endocrinol Metab 27, 84–95. 10.1016/j.tem.2015.11.008.

33. Caballero, F., Fernandez, A., De Lacy, A.M., Fernandez-Checa, J.C., Caballeria, J., and Garcia-Ruiz, C. (2009). Enhanced free cholesterol, SREBP-2 and StAR expression in human NASH. J Hepatol 50, 789–796. 10.1016/j.jhep.2008.12.016.

34. Behari, J., Yeh, T.H., Krauland, L., Otruba, W., Cieply, B., Hauth, B., Apte, U., Wu, T., Evans, R., and Monga, S.P. (2010). Liver-specific beta-catenin knockout mice exhibit defective bile acid and cholesterol homeostasis and increased susceptibility to diet-induced steatohepatitis. Am J Pathol 176, 744–753. 10.2353/ajpath.2010.090667.

35. Saponara, E., Penno, C., Orsini, V., Wang, Z.Y., Fischer, A., Aebi, A., Matadamas-Guzman, M.L., Brun, V., Fischer, B., Brousseau, M., et al. (2023). Loss of Hepatic Leucine-Rich Repeat-Containing G-Protein Coupled Receptors 4 and 5 Promotes Nonalcoholic Fatty Liver Disease. Am J Pathol 193, 161–181. 10.1016/j.ajpath.2022.10.008.

36. Yeh, T.H., Krauland, L., Singh, V., Zou, B., Devaraj, P., Stolz, D.B., Franks, J., Monga, S.P., Sasatomi, E., and Behari, J. (2010). Liver-specific beta-catenin knockout mice have bile canalicular abnormalities, bile secretory defect, and intrahepatic cholestasis. Hepatology 52, 1410–1419. 10.1002/hep.23801.

37. Pathak, P., and Chiang, J.Y.L. (2019). Sterol 12alpha-Hydroxylase Aggravates Dyslipidemia by Activating the Ceramide/mTORC1/SREBP-1C Pathway via FGF21 and FGF15. Gene Expr 19, 161–173. 10.3727/105221619X15529371970455.

38. Brufau, G., Stellaard, F., Prado, K., Bloks, V.W., Jonkers, E., Boverhof, R., Kuipers, F., and Murphy, E.J. (2010). Improved glycemic control with colesevelam treatment in patients with type 2 diabetes is not directly associated with changes in bile acid metabolism. Hepatology 52, 1455–1464. 10.1002/hep.23831.

39. Haeusler, R.A., Astiarraga, B., Camastra, S., Accili, D., and Ferrannini, E. (2013). Human insulin resistance is associated with increased plasma levels of 12alpha-hydroxylated bile acids. Diabetes 62, 4184–4191. 10.2337/db13-0639.

40. Bertaggia, E., Jensen, K.K., Castro-Perez, J., Xu, Y., Di Paolo, G., Chan, R.B., Wang, L., and Haeusler, R.A. (2017). Cyp8b1 ablation prevents Western diet-induced weight gain and hepatic steatosis because of impaired fat absorption. Am J Physiol Endocrinol Metab 313, E121–E133. 10.1152/ajpendo.00409.2016.

41. Kaur, A., Patankar, J.V., de Haan, W., Ruddle, P., Wijesekara, N., Groen, A.K., Verchere, C.B., Singaraja, R.R., and Hayden, M.R. (2015). Loss of Cyp8b1 improves glucose homeostasis by increasing GLP-1. Diabetes 64, 1168–1179. 10.2337/db14-0716.

42. Li, P., Ruan, X., Yang, L., Kiesewetter, K., Zhao, Y., Luo, H., Chen, Y., Gucek, M., Zhu, J., and Cao, H. (2015). A liver-enriched long non-coding RNA, lncLSTR, regulates systemic lipid metabolism in mice. Cell Metab 21, 455–467. 10.1016/j.cmet.2015.02.004.

43. Ma, B., and Hottiger, M.O. (2016). Crosstalk between Wnt/beta-Catenin and NF-kappaB Signaling Pathway during Inflammation. Front Immunol 7, 378. 10.3389/fimmu.2016.00378.

44. Nejak-Bowen, K., Kikuchi, A., and Monga, S.P. (2013). Beta-catenin-NF-kappaB interactions in murine hepatocytes: a complex to die for. Hepatology 57, 763–774. 10.1002/hep.26042.

45. Cobb, J.G., and Paschal, C.B. (2009). Improved in vivo measurement of myocardial transverse relaxation with 3 Tesla magnetic resonance imaging. J Magn Reson Imaging 30, 684–689. 10.1002/jmri.21877.

46. Lugert, S., Vogt, M., Tchorz, J.S., Muller, M., Giachino, C., and Taylor, V. (2012). Homeostatic neurogenesis in the adult hippocampus does not involve amplification of Ascl1(high) intermediate progenitors. Nat Commun 3, 670. 10.1038/ncomms1670.

47. Schuierer, S., Carbone, W., Knehr, J., Petitjean, V., Fernandez, A., Sultan, M., and Roma, G. (2017). A comprehensive assessment of RNA-seq protocols for degraded and low-quantity samples. BMC Genomics 18, 442. 10.1186/s12864-017-3827-y.

48. Liu, Z., Zhang, Y., Li, H., Guo, K., Tian, M., Cao, D., Kang, D.D., Xue, Y., Hou, X., Wang, C., et al. (2025). Furan-Derived Lipid Nanoparticles for Transporting mRNA to the Central Nervous System. Journal of the American Chemical Society. 10.1021/jacs.4c16326.

49. Leung, A.K.K., Hafez, I.M., Baoukina, S., Belliveau, N.M., Zhigaltsev, I.V., Afshinmanesh, E., Tieleman, D.P., Hansen, C.L., Hope, M.J., and Cullis, P.R. (2012). Lipid Nanoparticles Containing siRNA Synthesized by Microfluidic Mixing Exhibit an Electron-Dense Nanostructured Core. The Journal of Physical Chemistry C 116, 18440–18450. 10.1021/jp303267y.

50. Bligh, E.G., and Dyer, W.J. (1959). A rapid method of total lipid extraction and purification. Can J Biochem Physiol 37, 911–917. 10.1139/o59-099.

