## Supplementary material for "Targeting ZNRF3 and RNF43 to Restore Regeneration and Reverse Metabolic Dysfunction-Associated Steatotic Liver Disease": Suppl Figures and Legends

#### Supplemental information

#### Supplementary figure titles and legends

### Suppl Figure 1

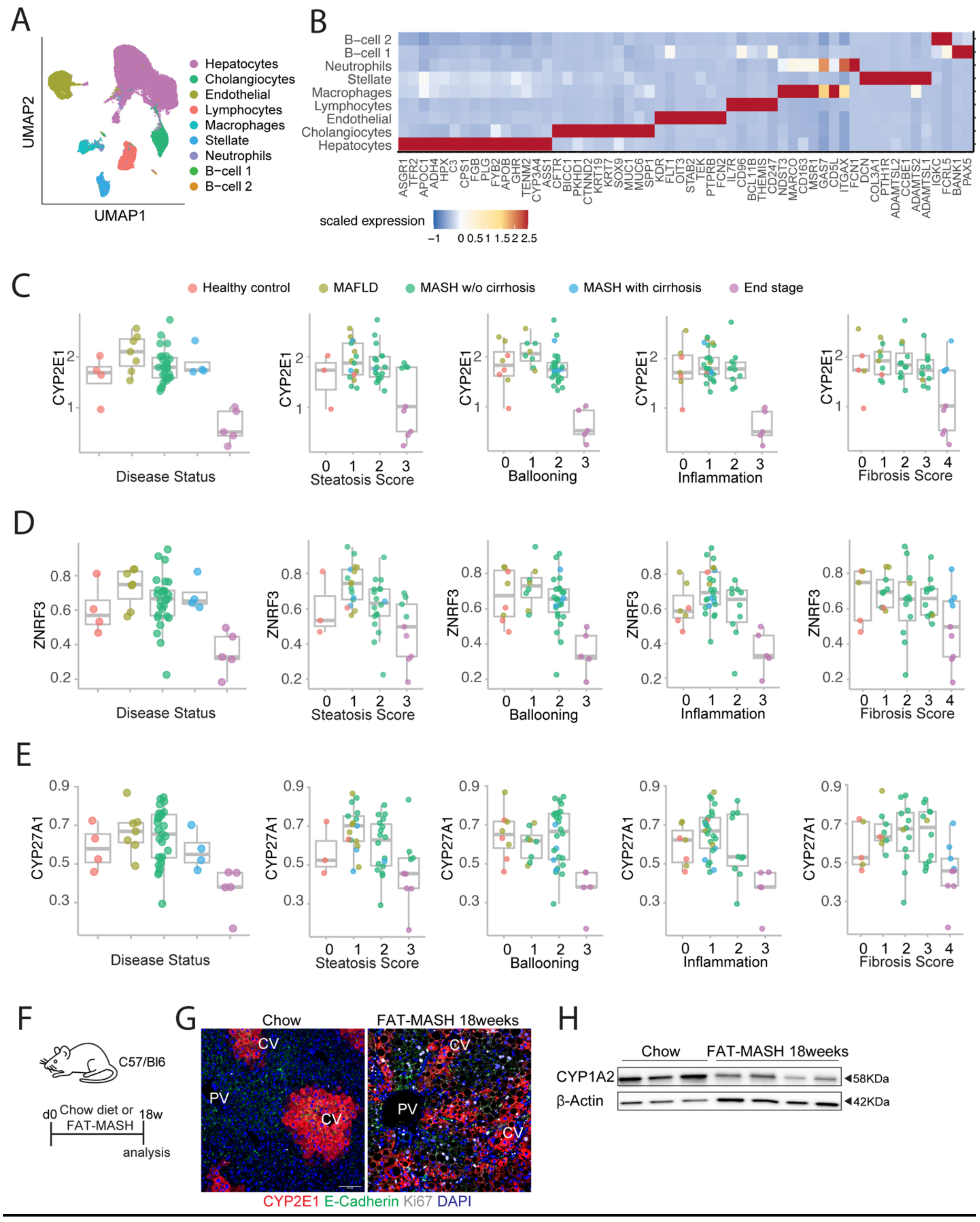

**<u>Supplementary Figure 1.</u> Integrated single-nucleus transcriptomics and correlation analyses for hepatocyte β-Catenin activity and MASLD/MASH progression, related to Figure 1**

(**A**) UMAP visualization of major liver cell populations human liver snRNA-seq data (GSE202379; n = 4 healthy, 7 MASLD, 27 MASH no cirrhosis, 4 MASH with cirrhosis, 5 end-stage). (**B**) Scaled expression heatmap of representative marker genes distinguishing the annotated cell types shown in (A). **(C–E)** Mean expression of individual β-Catenin pathway genes in hepatocytes, averaged per patient and stratified by clinical and histological features. Each dot represents one patient and is color-coded by disease stage. (C) CYP2E1, (D) ZNRF3, and (E) CYP27A1 expression levels plotted against disease status (left box plots) and key histological scores: steatosis, hepatocyte ballooning, lobular inflammation, and fibrosis stage (right four box plots). (**F**) Schematic of FAT-MASH model in ZNRF3/RNF43$^{fl/fl}$ mice used for G-H. (**G**) Representative IF staining for CYP2E1. (**H**) Western blot analysis of CYP1A2 and GAPDH.

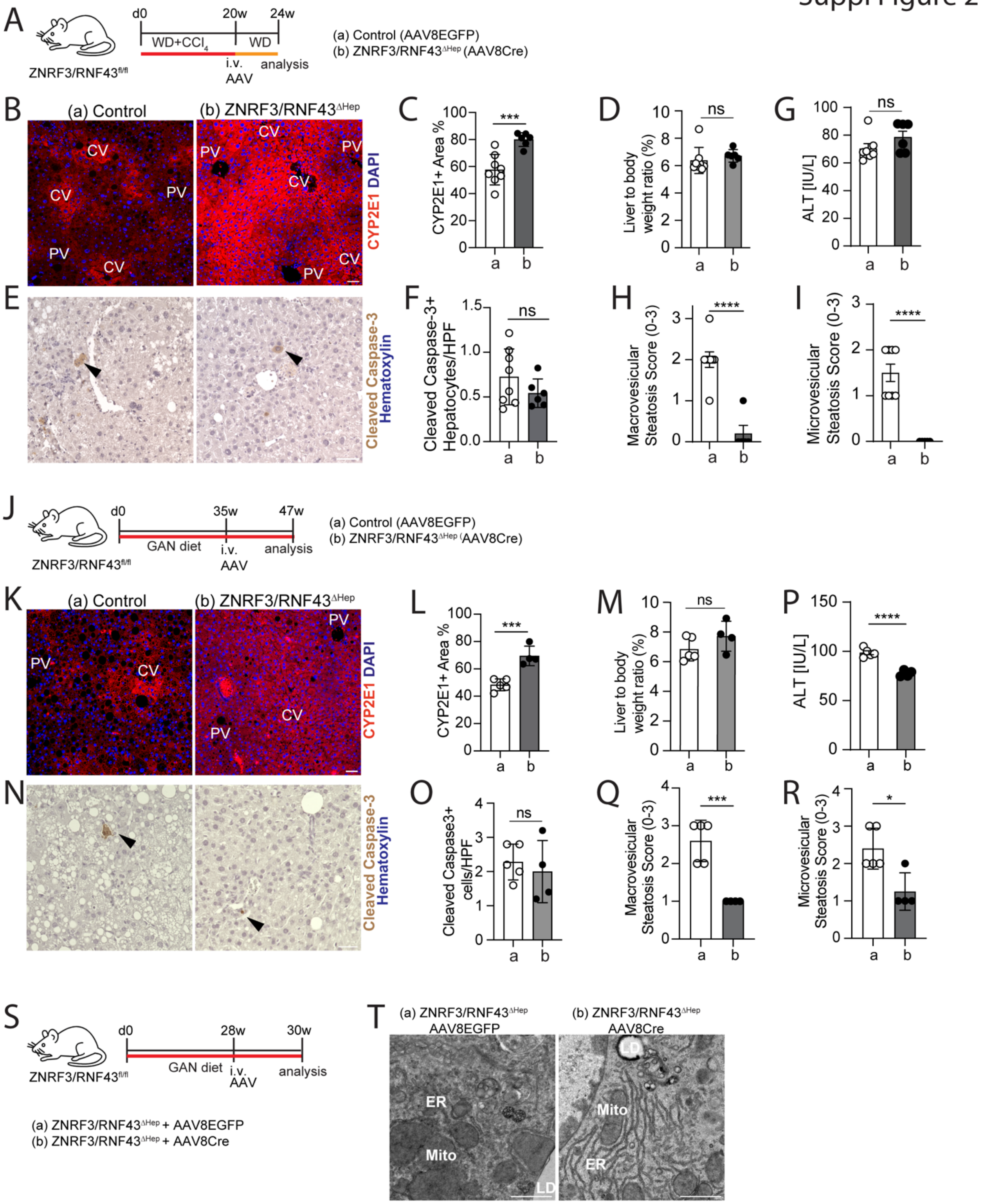


**Supplementary Figure 2.** **Histological and metabolic characterization of ZRdKO induced fatty liver disease regression, related to Figure 2.**

(**A)** Schematic of FAT-MASH model in ZNRF3/RNF43$^{fl/fl}$ mice used for B-I. **(B)** Representative IF staining for CYP2E1. **(C)** Quantification of CYP2E1-positive area (%) in liver sections. **(D)** Liver-to-body weight ratio (%). **(E)** Representative IHC images for cleaved Caspase-3. **(F)** Quantification of cleaved Caspase-3-positive hepatocytes per high-power field (HPF). **(G)** Serum ALT levels [IU/L]. **(H–I)** Histopathological scoring of macrovesicular (H) and microvesicular (I) steatosis. **(J)** Schematic of GAN diet-induced model in ZNRF3/RNF43$^{fl/fl}$ mice for J-R. **(K)** Representative IF staining for CYP2E1. **(L)** Quantification of CYP2E1-positive area (%) in liver sections. **(M)** Liver-to-body weight ratio (%). **(N)** Representative IHC for cleaved Caspase-3. **(O)** Quantification of cleaved Caspase-3-positive cells per HPF. **(P)** Serum ALT levels [IU/L]. **(Q–R)** Histopathological scoring of macrovesicular (Q) and microvesicular (R) steatosis. **(S)** Schematic of GAN diet-induced model in ZNRF3/RNF43$^{fl/fl}$ mice for T. **(T)** Representative electron microscopy (EM) images of hepatocytes from control and ZRdKO mice showing endoplasmic reticulum and mitochondrial structures. Data are presented as mean ± s.d. **with** individual mouse values overlaid as dots. Statistical significance was determined using a two-tailed unpaired Student's t-test. $p < 0.05$ (*), $p < 0.001$ (***), $p < 0.0001$ (****); ns, not significant. PV: portal vein; CV: central vein; ER: endoplasmic reticulum; Mito: mitochondrial. Scale bars: 100 µm (B, E, K, N), 1 µm (T),

Suppl Figure 3

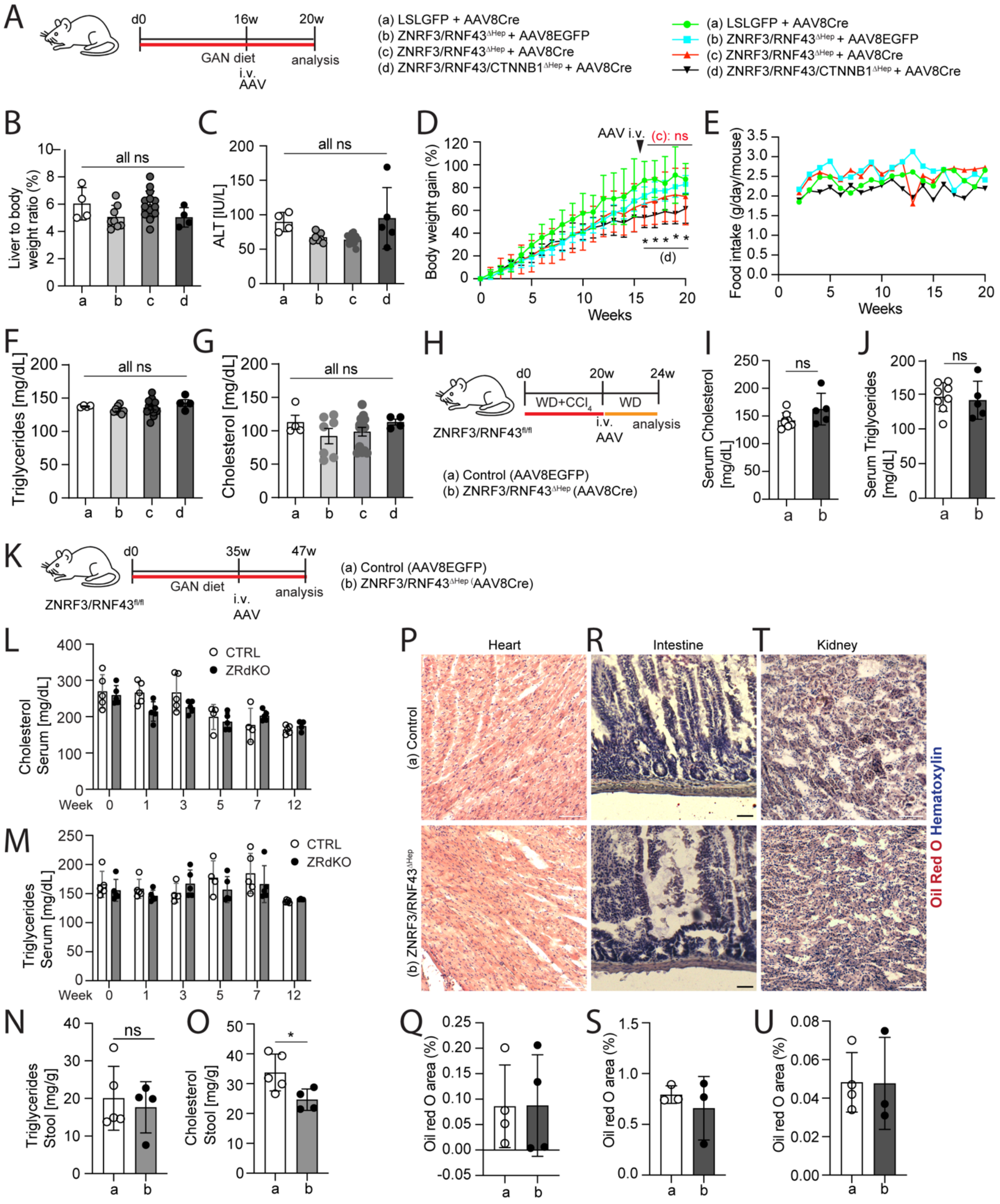


**Supplementary Figure 3. Histological and metabolic characterization of ZNRF3/RNF43 and ZNRF3/RNF43/CTNNB1 deletion mice in fatty liver diseases, related to Figure 3-4.**

(**A**) Schematic of the GAN diet-induced MASH model in four groups of mice for B-G: (a) R26$^{LSL\text{-}GFP}$ + AAV8-TBG-Cre (Cre-only control), (b) ZNRF3/RNF43$^{fl/fl}$ + AAV8-TBG-EGFP

(virus-only control), (c) ZNRF3/RNF43$^{\Delta hep}$ + AAV8-TBG-Cre (ZRdKO), (d) ZNRF3/RNF43/CTNNB1$^{\Delta hep}$ + AAV8-TBG-Cre (triple knockout). (**B**) Liver-to-body weight ratio (%). (**C**) Serum ALT levels [IU/L]. (**D**) longitudinal analysis of body weight gain (%). (**E**) food intake per gram per mouse per day. **(F–G)** Serum triglyceride and cholesterol concentrations. (**H**) Schematic of FAT-MASH model for ZNRF3/RNF43$^{fl/fl}$ mice received an intravenous injection of AAV8-EGFP (a, Control) or AAV8-Cre (b, ZNRF3/RNF43$^{\Delta hep}$). **(I–J)** Serum cholesterol and triglyceride levels. **(K)** Schematic of GAN diet-induced model in ZNRF3/RNF43$^{fl/fl}$ mice with the following treatment: (a) Control (AAV8-EGFP); (b) ZNRF3/RNF43$^{\Delta hep}$ (AAV8-Cre) used for L–U. **(L–M)** Bi-weekly measurements of serum cholesterol and triglycerides during regression. **(N–O)** Triglyceride and cholesterol levels in feces. (**P, R, T**) Representative Oil Red O images of heart (P), small intestine (R), and kidney (T) sections and their quantification respectively (**Q, S, U**).
Data are presented as mean ± s.d. with individual mouse values overlaid as dots. Statistical significance was determined using a two-tailed unpaired Student's t-test except for one-way ANOVA with multiple comparisons for panels B, C, F, and G. $p < 0.05$ (*); ns, not significant. Scale bars: 100 μm (P, R, T).

Suppl Figure 4

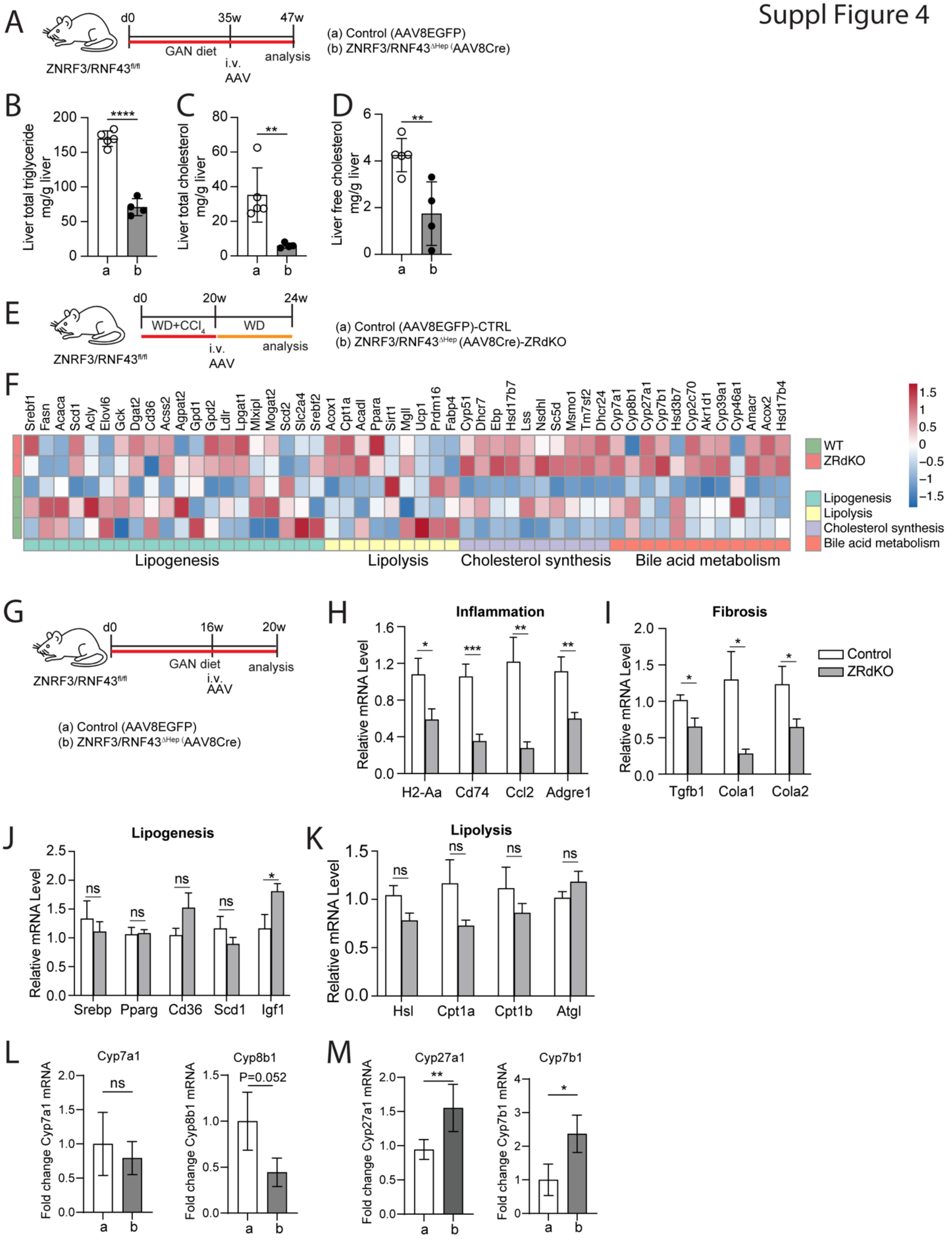

**<u>Supplementary Figure 4.</u> ZNRF3/RNF43 deletion leads to metabolic reprogramming, related to Figure 4.**

**(A)** Schematic of GAN diet-induced model in ZNRF3/RNF43$^{fl/fl}$ mice. Two experimental groups were generated: (a) Control: ZNRF3/RNF43$^{fl/fl}$ + AAV8-EGFP, (b) ZNRF3/RNF43$^{fl/fl}$ + AAV8-Cre. **(B–D)** Quantification of total liver triglycerides **(B)**, total liver cholesterol **(C)**, and liver free cholesterol **(D)**. (**E**) Schematic of FAT-MASH model in ZNRF3/RNF43$^{fl/fl}$ mice used for F. **(F)** Heatmap showing expression of selected genes involved in lipid metabolism pathways. **(G)** Schematic of GAN diet- induced model in ZNRF3/RNF43$^{fl/fl}$ mice used for H-K. **(H–K)** qPCR analysis of genes involved in inflammation (H), fibrosis (I), lipogenesis (J), and lipolysis (K). **(L)** qPCR analysis of classical bile acid synthesis genes *Cyp7a1* and *Cyp8b1*. **(M)** qPCR analysis of alternative bile acid synthesis genes *Cyp27a1* and *Cyp7b1*. Data are presented as mean ± s.d., except for qPCR panels (H–M), which are shown as mean ± s.e.m., with individual mouse values overlaid as dots. Statistical significance was determined using a two-tailed unpaired Student's t-test. $p < 0.05$ (*), $p < 0.01$ (**),$p < 0.0001$ (****); ns, not significant.

Suppl Figure 5

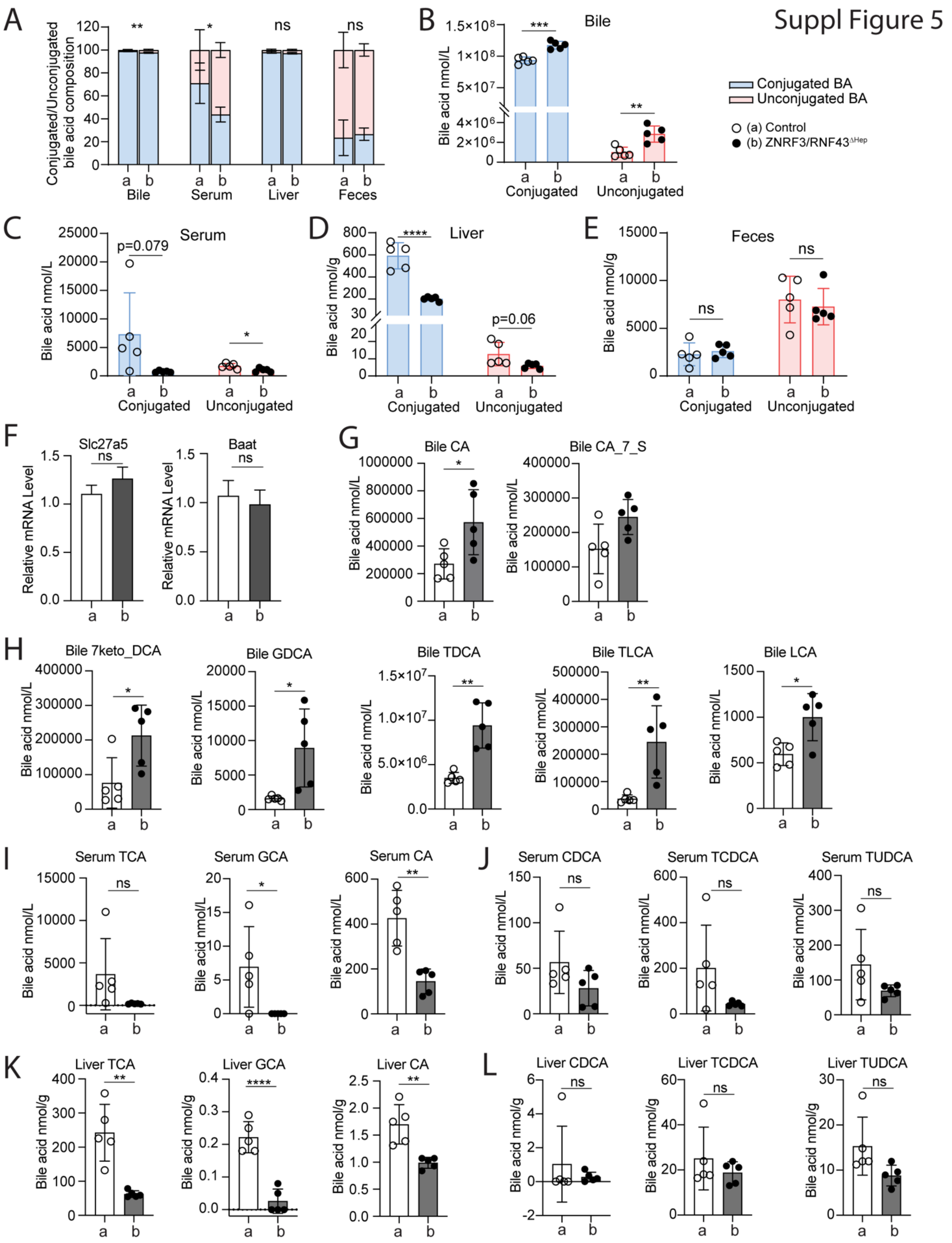

**Supplementary Figure 5. Comprehensive analysis of bile acid composition following ZNRF3/RNF43 deletion, related to Figure 5.**

**(A)** Stacked bar plots showing the proportion of conjugated (light blue) and unconjugated (light pink) bile acids in bile, liver, serum, and feces across groups (a) Control and (b) ZNRF3/RNF43$^{\Delta hep}$. **(B-E)** Total bile acid concentrations classified into conjugated or unconjugated bile acids in bile **(B),** serum (**C**), liver (**D**) and feces (**E**). (**F**) qPCR analysis of bile acid conjugation genes *Slc27a5* and *Baat*. **(G-L)** List of bile acids and their concentrations were shown as in the plot from bile (**G-H**), serum (**I-J**) and liver (**K-L**). Data are presented as mean ± s.e.m. for qPCR panels (F) and mean ± s.d. for the rest, with individual mouse values overlaid as dots. Statistical significance was assessed using a two-tailed unpaired Student's t-test. $p < 0.05$ (*), $p < 0.01$ (**), $p < 0.001$ (***), $p < 0.0001$ (****); ns, not significant.

Suppl Figure 6

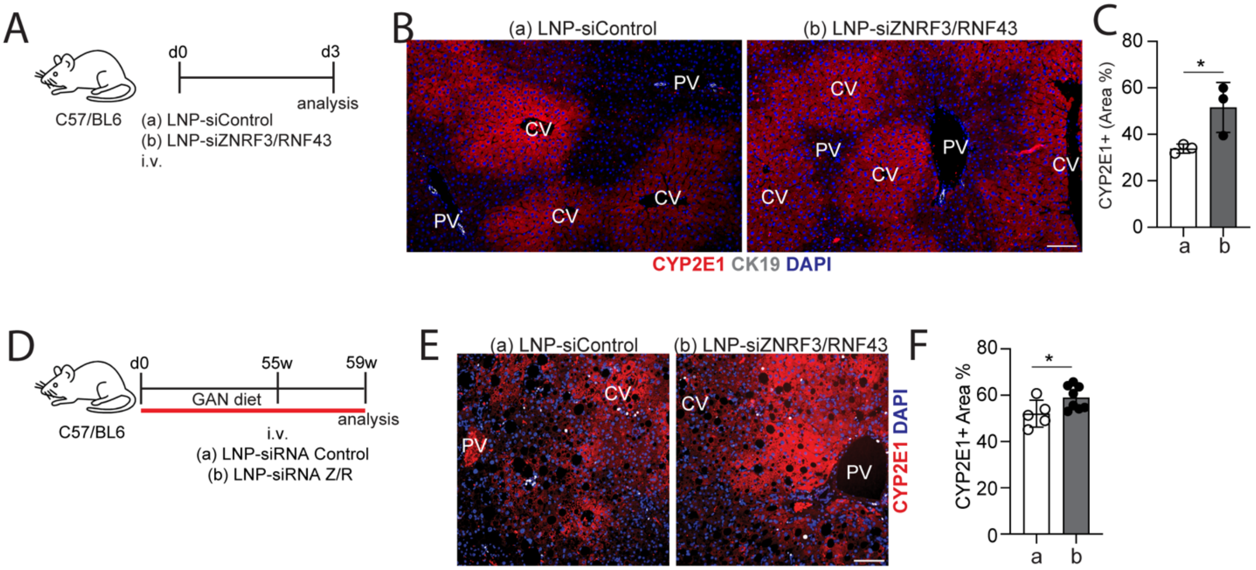


**Supplementary Figure 6. ZNRF3/RNF43 deletion serves as a therapeutic target, related to Figure 6**

**(A)** Schematic of LNP-siRNA delivery in WT C57BL/6J mice. (**B**) Representative IF staining for CYP2E1. (**C**) Quantification of CYP2E1-positive area (%). (**D**) Schematic of the GAN diet model used to test therapeutic LNP-siZNRF3/RNF43 intervention. **(E)** Representative CYP2E1 staining post-treatment. Data are presented as mean ± s.d., with individual mouse values overlaid as dots. PV: portal vein; CV: central vein. Statistical significance was assessed using a two-tailed unpaired Student's t-test. $p < 0.05$ (*). Scale bars: 100 μm (B, E)


1. Gribben, C., et al., *Acquisition of epithelial plasticity in human chronic liver disease.* Nature, 2024. **630**(8015): p. 166–173.

KEY RESOURCES TABLE

| Antibodies | | |
|---|---|---|
| **REAGENT or RESOURCE** | **SOURCE** | **IDENTIFIER** |
| Alexa Fluor–conjugated secondary antibody - donkey anti chicken – Alexa 488 | Jackson ImmunoResearch Milian Analytica AG | 703-545-155 |
| Alexa Fluor–conjugated secondary antibody - donkey anti goat – Alexa 647 | Jackson ImmunoResearch Milian Analytica AG | 705-605-147 |
| Alexa Fluor–conjugated secondary antibody - donkey anti mouse – Cy$^{TM}$3 | Jackson ImmunoResearch Milian Analytica AG | 715-165-151 |
| Alexa Fluor–conjugated secondary antibody - Donkey Anti-Rabbit – Alexa 647 | Jackson ImmunoResearch Milian Analytica AG | 711-605-152 |
| Alexa Fluor–conjugated secondary antibody - donkey anti rat – Cy$^{TM}$3 | Jackson ImmunoResearch Milian Analytica AG | 712-165-153 |
| Anti-mouse IgG-HRP | Bio-Rad | 1706516 |
| Anti-rabbit IgG-HRP | Cell Signaling Technology | 7074 |
| Chicken anti-Albumin | Sigma-Aldrich | SAB3500217 |
| Goat anti-GFP | Abcam | ab6673; RRID:AB_305643 |
| Histofine Simple Stain MAX PO anti-rabbit | Nichirei Biosciences | 414341F |
| Histofine Simple Stain MAX PO anti-rat | Nichirei Biosciences | 414311F |
| ImmPRESS anti-rabbit HRP reagent | Vector Laboratories | MP-7500 |
| Mouse anti-E-Cadherin | BD Biosciences | Cat# 610181; RRID:AB_397580 |
| Mouse anti-GAPDH | Proteintech | 60004-1-Ig |
| Rabbit anti-Cleaved Caspase-3 | Cell Signaling Technology | 9661S; RRID:AB_2341188 |
| Rabbit anti-CYP1A2 | Abcam | ab22717 |
| Rabbit anti-CYP27A1 | Abcam | ab126785 |
| Rabbit anti-CYP2E1 | Sigma-Aldrich | HPA009128 |
| Rabbit anti-CYP7B1 | Proteintech | 82927-1-RR |
| Rabbit anti-Ki67 | Abcam | ab15580 |
| Rat anti-CK19 | DSHB | TROMA-III; RRID:AB_2133570 |

| Chemicals | | |
|---|---|---|
| **REAGENT or RESOURCE** | **SOURCE** | **IDENTIFIER** |

| | | |
|---|---|---|
| Aprotinin | Thermo Scientific | 78432 |
| BSA | Bio-Rad | 500-0007 |
| Beta-mercaptoethanol | Sigma-Aldrich | 63689 |
| Chloroform | Sigma-Aldrich | C2432 |
| Citrate Buffer (10X, pH 6.0) | Sigma-Aldrich | C9999 |
| Coat-Quick Glo adhesive pen | Electron Microscopy Sciences | 10310 |
| D-Luciferin, Potassium Salt | Thermo Scientific | 88292 |
| DAPI | Sigma-Aldrich | D9542 |
| Direct Red 80 | Sigma-Aldrich | 365548-5G |
| EDTA solution, 0.5 M, pH 8.0 (DNase-, RNase-, Protease-free) | Sigma-Aldrich | 03690 |
| Eosin Y | Sigma-Aldrich | HT110116 |
| Epon resin | Electron Microscopy Sciences | 14120 |
| Ethanol, absolute | Sigma-Aldrich | E7023 |
| Fast Green FCF | Sigma-Aldrich | F7258 |
| Formalin (10% NBF) | Thermo Fisher Scientific | 23-245685 |
| Glutaraldehyde (25%) | Electron Microscopy Sciences | 16220 |
| Glycerol | Sigma-Aldrich | G5516 |
| Halt,Ñ¢ Protease Inhibitor Cocktail | Thermo Scientific | 78429 |
| Hematoxylin | Sigma-Aldrich | H3136 |
| Hydrogen Peroxide | ACD | part of Cat# 323100 |
| ImmPACT DAB substrate kit | Vector Laboratories | SK-4105 |
| Isopropanol | Sigma-Aldrich | I9516 |
| Kimwipe | Kimberly-Clark | 34155 |
| Laemmli Sample Buffer | Bio-Rad | 1610737 |
| Lead citrate (Reynolds) | Electron Microscopy Sciences | 17800 |
| Leupeptin | Thermo Scientific | 78435 |
| Methanol | Sigma-Aldrich | 179337-4L |
| NaOH | Sigma-Aldrich | 34120 |
| Oil Red O powder | Sigma-Aldrich | O0625-100G |
| Osmium tetroxide (4%) | Electron Microscopy Sciences | 19100 |
| PBS | Thermo Fisher Scientific | 10010-023 |
| PMSF | Sigma-Aldrich | P7626-1G |
| Paraformaldehyde (PFA) powder | Sigma-Aldrich | P6148 |
| Pepstatin A | Thermo Scientific | 78436 |
| Permount | Fisher Scientific | SP15-100 |
| Picric acid | Sigma-Aldrich | P6744 |
| Propylene oxide | Electron Microscopy Sciences | 20401 |
| Proteinase K | Sigma-Aldrich | P2308 |
| Reagent A and B (BCA Kit) | Thermo Fisher Scientific | 23225 |

| | | |
|---|---|---|
| SYBR Green Supermix | Bio-Rad | 1725121 |
| Sodium cacodylate buffer (0.2 M, pH 7.2) | Electron Microscopy Sciences | 11652 |
| Sodium phosphate buffer (0.1 M, pH 7.2) | Thermo Fisher Scientific | 1869126 |
| Toluidine Blue O | Sigma-Aldrich | T3260 |
| Triton X-100 | Sigma-Aldrich | T8787-250ML |
| Tween-20 | Sigma-Aldrich | P9416 |
| Uranyl acetate | Electron Microscopy Sciences | 22400 |
| Xylene | Thermo Fisher Scientific | HC7001GAL |
| Cholesterol E | FUJIFILM | Cat# 999-02601 |
| Free Cholesterol E | FUJIFILM | Cat# 993-02501 |
| L-Type Triglyceride M Enzyme Color A | FUJIFILM | Cat# 994-02891 |
| L-Type Triglyceride M Enzyme Color B | FUJIFILM | Cat# 990-02991 |
| Multi-Calibrator Lipid | FUJIFILM | Cat# 464-01601 |

| **Critical Commercial Assays** | | |
|---|---|---|
| **REAGENT or RESOURCE** | **SOURCE** | **IDENTIFIER** |
| ALT (GPT) Reagent Set | Teco Diagnostics | A524-150 |
| Cholesterol Reagent Kit | Teco Diagnostics | C518-120 |
| High-Capacity cDNA Reverse Transcription Kit | Thermo Fisher Scientific | 4368813 |
| Pierce BCA Protein Assay Kit | Thermo Fisher Scientific | 23225 |
| RNAscope Hydrogen Peroxide | ACD | part of Cat# 323100 |
| RNAscope Multiplex Fluorescent Reagent Kit v2 | ACD | 323100 |
| RNAscope Protease Plus | ACD | part of Cat# 323100 |
| RNAscope Target Retrieval Reagent | ACD | part of Cat# 323100 |
| RNeasy Plus Mini Kit | QIAGEN | 74134 |
| Triglyceride GPO Liquid Reagent Kit | Teco Diagnostics | T7532-150 |
| TruSeq RNA Sample Prep Kit v2 | Illumina | RS-122-2001 |

| **Deposited Data** | | |
|---|---|---|
| **REAGENT or RESOURCE** | **SOURCE** | **IDENTIFIER** |
| Bulk RNA-seq data (ZNRF3/RNF43 WT and ΔHep) | NIH Sequence Read Archive | BioProject ID: XXXXXXXX |

| **Dietary Regimens** | | |
|---|---|---|
| **REAGENT or RESOURCE** | **SOURCE** | **IDENTIFIER** |

| | | |
|---|---|---|
| FAT-MASH Diet ("Western"/Fast Food Diet) - 20-23% Fat by weight (40-45% kcal from Fat), Milkfat (SFA >60% of total Fatty Acids), 1.25% Cholesterol | Envigo | TD.120528 |
| GEN Diet - Rodent Diet With 40 kcal% Fat (Palm Oil), 20 kcal% Fructose and 2% Cholesterol | Research Diets Inc. | D09100310 |

| **Equipment** | |
|---|---|
| **REAGENT or RESOURCE** | **SOURCE** |
| Advantage CCD Camera | Advanced Microscopy Techniques |
| Hitachi 7700 TEM | Hitachi High-Technologies Corp. |
| Leica UCT Ultramicrotome | Leica Microsystems |
| Stand Axio Observer 7 Microscope | Carl Zeiss AG |

| **Experimental Models: Organisms/Strains** | |
|---|---|
| **REAGENT or RESOURCE** | **SOURCE** |
| C57BL/6 | Jackson Laboratory |
| R26-STOP-EGFP | Novartis |
| ZNRF3/RNF43$^{Flox/Flox}$ | Novartis |
| ZNRF3/RNF43/CTNNB1$^{\Delta Hep}$ | This study |

| **Oligonucleotides (Primers)** | | |
|---|---|---|
| **GENE** | **FORWARD PRIMER** | **REVERSE PRIMER** |
| Abca1 | TCCTGATCTCTGTACGCCTGAGCTACCC | GAGTTGGATAACGGAAGCAGGGGTTGTTGGC |
| Abcg5 | CAGCTTAGGTGTCCTGCATGTGTC | CAAGGAGACATCTTTGAGGATTTGCC |
| Abcg8 | TCAGAGAGACCCTGGCTTTCATTG | GCAGCTCGGCGATTACGTCTTC |
| Apoa1 | CAGAGACTATGTGTCCCAGTTTGAATCCT | TTATCCCAGAAGTCCCGAGTCAATGGGCCC |
| Atgl | GGATGGCGGCATTTCA | CAAAGGGTTGGGTTGG |
| Baat | GGATAGCCTGACTCTGGAAAGG | CAATCCACCAGCACCTCCAAAC |
| Abca11 | TCTGACTCAGTGATTCTTCGCA | CCCATAAACATCAGCCAGTTGT |
| CD74 | AGTGCGACGAGAACGGTAAC | CGTTGGGGAACACACACCA |
| Ccl2 | AGGTCCCTGTCATGCTTCTG | TCTGGACCCATTCCTTCTTG |
| Cd36 | AAAACGACTGCAGGTCAACA | GCAACAAACATCACCACTCC |
| Cpt1a | CCTGGGCATGATTGCAAAG | GGACGCCACTCACGATGTT |
| Cpt1b | GGCTGCCGTGGGACATT | TGCCTTGGCTACTTGGTACGA |
| H2Aa | TCAGTCGCAGACGGTGTTTAT | GGGGGCTGGAATCTCAGGT |
| Hsl | GAGCGCTGGAGGAGTGTTTT | TGATGCAGAGATTCCCACCTG |
| Igf1 | TGGTGGATGCTCTTCAGTTC | TGAGTCTTGGGCATGTCAGT |

| | | |
|---|---|---|
| Mttp | GATGTGGACGTTGTGTTACTGTG GAGGAATC | GAAGATGCTCTTCTCGCCTCTCT GTTGAC |
| Slc10a1 | CAAACCTCAGAAGGACCAAACA | GTAGGAGGATTATTCCCGTTGTG |
| Pparg | GACCAGGGAGTTCCTCAAAA | CAGCAGGTTGTCTTGGATGT |
| Scd1 | AGCCTGTTCGTTAGCACCTT | GGGAAGGTGTGGTGGTAGTT |
| Shp | CCAAGGAGTATGCGTACCTGAAG | GCTCCAAGACTTCACACAGTGC |
| Slc27a5 | CTGCGGTACTTGTGTAACGTCC | TCCGAATGGGACCAAAGCGTTG |
| Srebp1c | GGAGCCATGGATTGCACATT | GGCCCGGGAAGTCACTGT |
| Tgfb1 | CTCCCGTGGCTTCTAGTGC | GCCTTAGTTTGGACAGGATCTG |

| **DsiRNA** | | |
|---|---|---|
| **REAGENT or RESOURCE** | **Sense** | **Anti-sense** |
| *mm.Ri.Znrf3.13.1* | *mCmArC mArUmU rUrGrA rGrAmU rGmArU mCrAmG rArArU rArCmU T* | *rArAmG rUrArU rUrCmU rGmArU mCrArU rCrUrC rArArA rUmGrU mGmCmA* |
| *mm.Ri.Rnf43.13.1* | *mGmCrA mArCmU rUrCrA rGrCmU rAmUrA mUrCmA rUrUrU rCrCmU G* | *rCrAmG rGrArA rArUmG rAmUrA mUrArG rCrUrG rArArG rUmUrG mCmCmC* |
| *Negative control* | *mCmGrUmUrAmArUrCrGr CrGmUrAmUrAmArUmArCr GrCrGrUmAT* | *rArUmArCrGrCrGrUmArUm UrAmUrArCrGrCrGrArUrUr AmArCmGmAmC* |
| **AAV virus** | | |
| **REAGENT or RESOURCE** | **SOURCE** | **IDENTIFIER** |
| AAV8.TBG.PI.Cre.rBG | Addgene | Cat# 107787-AAV8 |
| pAAV.TBG.PI.eGFP.WPRE .bGH | Addgene | Cat# 105535-AAV8 |

| **Software and Algorithms** | | |
|---|---|---|
| **REAGENT or RESOURCE** | **SOURCE** | **IDENTIFIER / CITATION** |
| DESeq2 v1.42.0 | Bioconductor | Love MI, Huber W, Anders S (2014). Moderated estimation of fold change and dispersion for RNA-seq data with DESeq2. Genome Biology, 15(12), 550. DOI: https://doi.org/10.1186/s13059-014-0550-8 |
| ggplot2 | CRAN / Tidyverse | Wickham H (2016). ggplot2: Elegant Graphics for Data Analysis. Springer-Verlag New York. ISBN: 978-3-319-24277-4. URL: https://ggplot2.tidyverse.org |
| GraphPad Prism v10 | GraphPad Software | RRID:SCR_002798 |
| ImageJ (Fiji) | NIH | RRID:SCR_003070. URL: https://imagej.net |
| LabSolutions Insight | Shimadzu Corporation | https://www.shimadzu.com/an/products/liquid-chromatograph-mass-spectrometry/lc-ms-software/labsolutions-insight/index.html |

| | | |
|---|---|---|
| Living Image Software | Caliper Life Sciences | RRID:SCR_014247. Purchased with the support of NCRR S10-RR026561-01 |
| R v4.4.1 | R Foundation for Statistical Computing | R Core Team (2025). R: A language and environment for statistical computing. URL: https://www.R-project.org/ |
| Seurat v5.1.0 | Satija Lab / CRAN | Hao Y, Stuart T, Kowalski MH, et al. (2023). Dictionary learning for integrative, multimodal and scalable single-cell analysis. Nature Biotechnology. DOI: https://doi.org/10.1038/s41587-023-01767-y |
| STAR v2.7.11a | Dobin et al. | Dobin A, Davis CA, Schlesinger F, et al. (2013). STAR: ultrafast universal RNA-seq aligner. Bioinformatics, 29(1), 15–21. DOI: https://doi.org/10.1093/bioinformatics/bts635 |
